# Heat-killed Latilactobacillus sakei D279 Attenuates Lipopolysaccharide-Induced Intestinal Inflammation and Barrier Disruption

**DOI:** 10.64898/2026.11.10.765754

**Authors:** Ryota Chihama, Takamasa Tsunoda, Nur Syafiqah Mohamad Ishak, Katsuyuki Hashimoto

**Affiliations:** Niigata Research Laboratory, Mitsubishi Gas Chemical Company, Inc., 182 Tayuhama, Kita-ku, Niigata 950-3112, Japan

**Keywords:** *Latilactobacillus sakei*, lipopolysaccharide, intestinal barrier, inflammation, MLCK, gut microbiota, postbiotic

## Abstract

Intestinal barrier dysfunction and excessive inflammation contribute to the pathogenesis of various metabolic and gastrointestinal diseases. Lipopolysaccharide (LPS) induces systemic and intestinal inflammation, disrupts epithelial tight junctions, and impairs intestinal barrier integrity. Although heat-killed *Latilactobacillus sakei* D279 exhibits antiviral activity, its effects on intestinal barrier function remain unclear. In this study, we examined whether heat-killed D279 preserves intestinal barrier integrity and attenuates LPS-induced inflammatory responses in vivo (LPS-challenged mice), ex vivo (mouse peritoneal macrophages), and in vitro (IEC-6 intestinal epithelial cells). Cytokine and receptor gene expression in macrophages was measured to evaluate inflammatory responses. Furthermore, barrier function was assessed using transepithelial electrical resistance (TEER), and the expression of tight junction-related genes was assessed. D279 alleviated LPS-induced intestinal injury in LPS-challenged mice, as indicated by the attenuation of colon shortening and histopathological changes compared to those in the LPS control group. In peritoneal macrophages, D279 pretreatment suppressed LPS-induced expression of pro-inflammatory cytokines (*TNF-*α, *IL-1*β, and *IL-12*β), increased *IL-10 expression*, and reduced LPS-induced upregulation of *MLCK* expression. This postbiotic increased basal cell viability in a concentration-dependent manner in IEC-6 cells. Moreover, it enhanced epithelial barrier development (increased TEER) and upregulated tight junction gene expression. These findings demonstrate that heat-killed D279 primarily confers its benefits by attenuating LPS-induced cytokine responses, limiting the upregulation of *MLCK* expression, and partially preserving goblet cell abundance. This is consistent with its protective postbiotic role in intestinal homeostasis. Thus, this study introduces a candidate postbiotic for the promotion of intestinal resilience under systemic inflammatory stress.

## 1. Introduction

The intestinal barrier preserves gut health and maintains physiological homeostasis, serving serves as a selective interface between the intestinal lumen and host. Specifically, this barrier facilitates nutrient absorption while preventing the entry of pathogens, toxins, and microbial products [1,2]. The intestinal barrier is not a single layer; it is a multilayered defense system with multiple physical, chemical, microbiological, and immunological components that synergistically regulate the interactions between intestinal contents and the host [3]. The physical barrier is primarily formed by intestinal epithelial cells and tight junction (TJ) proteins, whereas the chemical barrier includes a mucus layer and antimicrobial factors that limit direct contact between luminal microorganisms and the intestinal epithelium. The intestinal microbiota and their metabolites maintain the microbiological barrier, whereas the immunological barrier consists of innate and adaptive immune mechanisms that regulate host responses to luminal stimuli. The coordinated regulation of these components is essential for maintaining intestinal homeostasis. Conversely, disruption of this integrated barrier increases intestinal permeability and permits the passage of luminal substances into the host body. Importantly, intestinal barrier dysfunction may have consequences beyond the intestinal tract because increased permeability can also contribute to systemic inflammatory and metabolic disturbances [3]. Thus, maintaining intestinal barrier integrity is important not only for local intestinal homeostasis but also for the overall preservation of physiological health.

Lipopolysaccharide (LPS), a toxin derived from the outer membrane of gram-negative bacteria, triggers inflammatory responses and compromises the host intestinal barrier. Excessive exposure to LPS can impair epithelial integrity by disrupting TJ proteins and stimulating the production of proinflammatory cytokines, thereby increasing intestinal permeability and promoting intestinal inflammation [4,5]. Persistent or excessive inflammatory responses further aggravate epithelial barrier damage and create a vicious cycle of inflammation and barrier dysfunction. Consequently, nutritional strategies that preserve the integrity of the intestinal barrier while appropriately modulating excessive inflammatory responses have garnered increasing attention as potential approaches for maintaining gut health and preventing intestinal barrier dysfunction[1,2].

Postbiotics, formulations consisting of non-living microorganisms and/or their components, provide health benefits to the host [6]. They have emerged as promising alternatives to live probiotics because of their favorable stability, safety, and ease of formulation [7,8]. Heat-killed lactic acid bacteria (LAB) retain biologically active cellular components that interact with host cells and modulate their physiological functions. For example, a heat-killed *Lacticaseibacillus paracasei* strain improved epithelial barrier function by restoring transepithelial electrical resistance (TEER) and increasing the expression of TJ proteins under inflammatory conditions[9]. In addition to their direct effects on epithelial cells, postbiotics may affect intestinal homeostasis by modulating immune responses and interacting with intestinal microbiota [10]. These properties suggest that heat-killed LAB contribute to the maintenance of intestinal barrier function through their complementary effects on the epithelial, immune, and microbiological components of the intestinal barrier.

We previously screened more than 1,000 LAB isolates with a specific focus on immune function to identify LAB strains with strong immunomodulatory potential. This screening revealed the strong immunomodulatory potential of *Latilactobacillus sakei* D279. D279 was originally isolated from traditional Japanese red turnip pickles (akakabu-zuke), and its heat-killed preparation has been commercially developed as “Liveole lactic acid bacteria.” Liveole LAB enhanced immune function and protected the host against virus-induced damage by preferentially inducing type III interferon (IFN-λ) and enhancing natural killer cell activity in mice infected with influenza [11]. These findings indicate that D279 possesses distinct immunomodulatory properties. However, whether heat-killed LAB can protect the intestinal barrier during systemic inflammatory stress and the epithelial and immune mechanisms involved remain unclear. To the best of our knowledge, no studies have established whether heat-killed *L. sakei* can preserve colonic barrier integrity in an intraperitoneal (i.p.) LPS challenge model mimicking systemic inflammation-driven intestinal injury.

In this study, we investigated the effects of heat-killed *L. sakei* D279 on intestinal inflammation and barrier integrity in an LPS-induced mouse model. Our primary objectives were to: i) characterize the physiological effects of systemic LPS challenge; ii) determine whether D279-mediated protection against secondary intestinal injury was accompanied by changes in the systemic biochemical status; and iii) explore the potential immunomodulatory mechanisms underlying these in vivo effects. By using this complementary approach, we aimed to determine whether heat-killed D279 protects intestinal homeostasis through the coordinated modulation of immune responses and epithelial barrier function, in addition to its effects on the mucus and intestinal microbial environment.

## 2. Materials and Methods

### 2.1. Preparation of Heat-Killed *Lactobacillus* and Reagents

*L. sakei* strain D279 (LiveOle) was grown in MRS broth at 37 °C for 24 h. Cells were collected via centrifugation at 10,000 × *g* for 5 min, and the culture supernatant was discarded. The bacterial pellet was washed via resuspension in distilled water and centrifuged at 10,000 × *g* for 5 min. After the supernatant was discarded, cells were resuspended in distilled water and subjected to heat treatment at 80 °C for 90 min, resulting in heat-killed bacteria. The resulting suspension was spray-dried to produce a heat-killed *L. sakei* D279 powder for storage and subsequent experiments. The inflammatory response was triggered in vitro, ex vivo, and in vivo using LPS derived from *Escherichia coli* O111:B4 (Sigma-Aldrich, St. Louis, MO, USA).

### 2.2 In Vivo Animal Experiment

#### 2.2.1. Animals and Housing Conditions

The animal experiments were approved by the Animal Care and Experimentation Facility Committee. The study was conducted in accordance with the committee’s rules. Four-week-old male C57BL/6J mice were obtained from The Jackson Laboratory (Yokohama, Japan). Animals (three per cage) were allowed to acclimate for 23 days. Housing conditions were controlled (temperature: 20– 26 °C; relative humidity: 40–70%; 12-h light/dark cycle with lights on from 07:00 to 19:00). The mice had continuous access to food and water and were housed in polyolefin cages that included bedding and enrichment throughout the study.

#### 2.2.2. Diets and Experimental Design

Mice were assigned to three groups (n = 6 per group, two cages per group): (i) control, (ii) LPS, and (iii) LPS+D279. During acclimation, the mice were fed a standard MF solid diet (Figure 1A). One week before heat-killed LAB feeding (day 0), the diet was switched to MF powder. From Day 8 onward, the Control and LPS groups continued to receive the MF powder diet, whereas the LPS+D279 group received the MF powder diet mixed with 0.5% (w/w) LAB (D279 strain).

**Figure 1:**
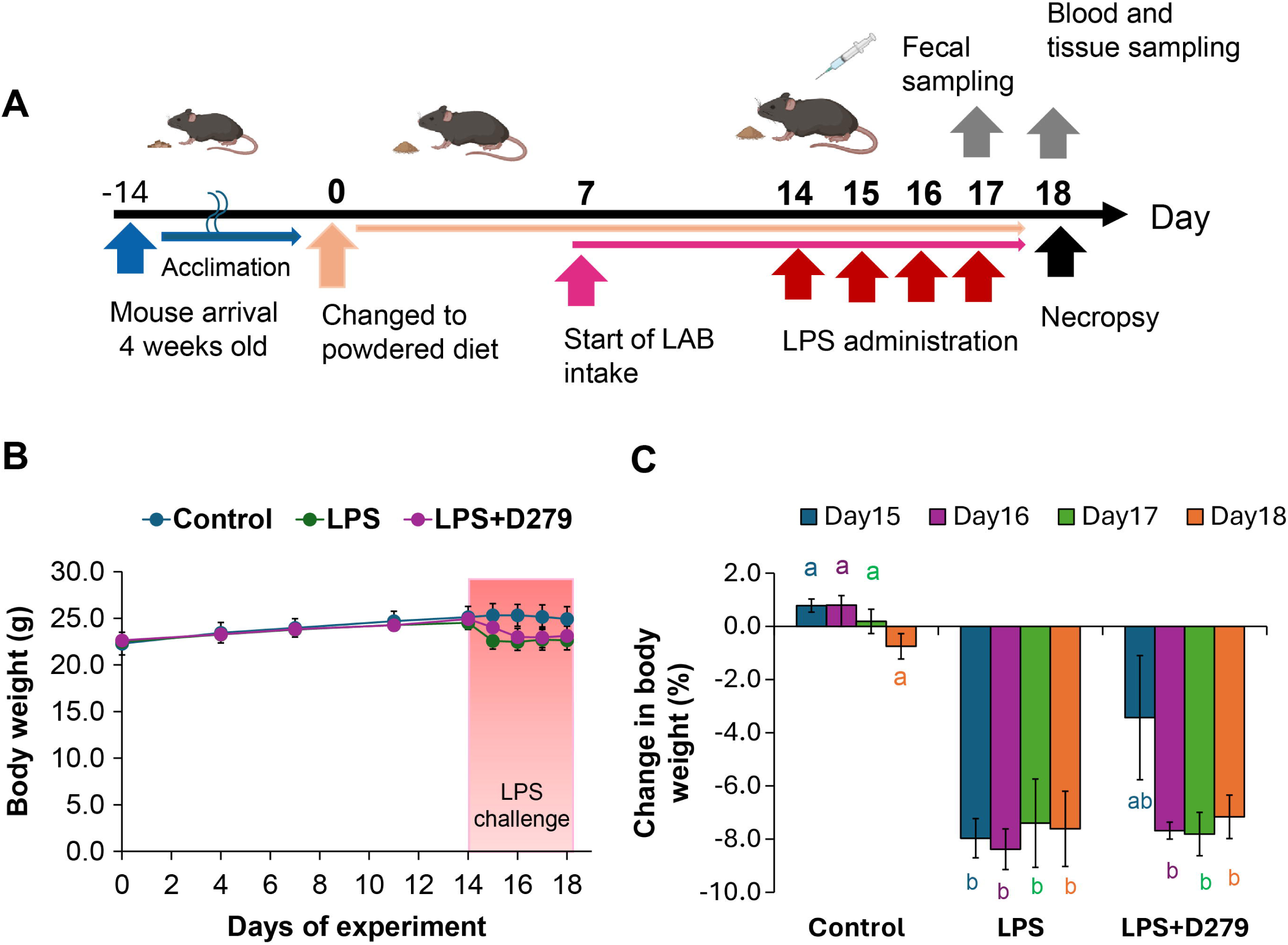
Effects of D279 supplementation and LPS challenge on body weight. (A) Experimental timeline showing the D279 supplementation period and subsequent LPS challenge. (B) Body weight during the experimental period. (C) Percentage change in body weight following LPS challenge, normalized to the pre-LPS challenge value (Day 14). Data are presented as the mean ± SD. Bars with different letters indicate significant differences among groups on the same day (one-way ANOVA followed by all pairwise multiple comparisons using the Holm–Sidak method; *p* < 0.05). Days 15–18 corresponded to 1–4 days post-LPS challenge. ANOVA, analysis of variance; LPS, lipopolysaccharide

#### 2.2.3. LPS Challenge

One week after initiating LAB feeding, intestinal barrier disruption was induced through the i.p. administration of LPS. LPS derived from *E. coli* O111:B4 was dissolved in phosphate-buffered saline (PBS) (pH 7.4; Gibco, Waltham, MA, USA) to prepare a 1.0 mg/mL stock solution, which was diluted to 0.1 mg/mL before dosing. The LPS and LPS+D279 groups were administered LPS (1.0 mg/kg) once daily for four consecutive days. The dosing volume was adjusted individually based on body weight using the following formula: body weight (g) × 0.01 (mL). The Control group received PBS only.

#### 2.2.4. Body Weight and Food Intake Monitoring

Body weight was measured twice weekly, starting on day 0 (Mondays and Fridays). Food intake was assessed twice weekly for each cage after switching to a powdered diet. This was achieved by weighing the feeders before and after use and correcting for spillage weight. The mean daily intake was calculated for each mouse.

#### 2.2.5. Necropsy and Sample Collection

Fresh fecal pellets were collected immediately after defecation the day before necropsy (day 13). Samples were stored at −30 °C for several hours and then transferred to −80 °C until further analyses (e.g., microbiota and triglyceride-related assays).

At the end of the study (day 14), the mice were euthanized under isoflurane inhalation anesthesia, and blood was collected from the abdominal vena cava of each mouse using a syringe. Whole blood (0.4–0.6 mL) was collected into BD Microtainer blood collection tubes containing a plasma separator and lithium heparin (DB Microtainer; 365985, Becton Dickinson, Japan) and mixed by gentle inversion (10 times). Samples were stored on ice until the completion of necropsy (up to approximately 2 h) and subsequently centrifuged at 6000 × *g* for 90 s at 4 °C to separate the plasma. Plasma was aliquoted into SUMILON serum tubes (1.0 mL; outer cap; Sumitomo Bakelite, Tokyo, Japan) and stored at −80 °C until use.

After euthanasia via exsanguination, the gastrointestinal tract was excised from the stomach to the anus, and the distance from the cecum to the anus was measured using a scale. The colon was defined as the proximal 2 cm from the cecum. The cecal-side proximal two-thirds was collected for tissue (histological) analysis, and the anal-side portion was collected for gene expression analysis. Similarly, the ileum was defined as the proximal 2 cm to the small intestine from the cecum and collected for subsequent analyses.

#### 2.2.6. Colon Histology and Imaging analyses

The colon specimens were immersed in 4% paraformaldehyde prepared in phosphate buffer and fixed for at least 24 h. Next, the tissues were routinely processed, embedded in paraffin, and sectioned for histological analysis. Hematoxylin and eosin (H&E) and Alcian blue–periodic acid– Schiff (AB–PAS) staining were performed by [excluded to facilitate double anonymization]. The stained slides were examined microscopically, and representative images were captured using a light microscope (Eclipse TC2000-S; Nikon, Tokyo, Japan) fitted with a digital camera (Ds-Ri2; Nikon).

Muscle layer thickness and villus length were quantified from H&E-stained images using the ImageJ software (National Institutes of Health, Bethesda, MD, USA). Measurements were performed on images from three colon sections. The resulting values were averaged per mouse and summarized as the mean ± standard error of the mean (SEM) for each group.

Goblet cells were evaluated in the AB-PAS-stained sections using a custom HTML-based image analysis application. A region of interest was defined for each image, and the area of Alcian blue– positive (blue-stained) cells within the region of interest was quantified. The resulting values were averaged per mouse and summarized as the mean ± SEM for each group. The HTML application code was generated with assistance from Exabase and reviewed and edited by the authors before use.

#### 2.2.7. Gene Expression Analysis

Samples designated for RNA isolation were placed in DNA/RNA Shield (Zymo Research, Irvine, CA, USA) in 2.0 mL tubes (TOMY SEIKO, Tokyo, Japan) and stored at −30 °C until processing. For tissue disruption, five to eight silicone beads (TOMY SEIKO) were added to each tube, followed by incubation with proteinase K at 25 °C for 1 h. The samples were then homogenized using a Micro Smash MS-100 Cell Disruptor (TOMY SEIKO). Total RNA was purified using a Quick-RNA Miniprep kit (Zymo Research) according to the manufacturer’s instructions.

Purified RNA was diluted to the required working concentration with DNase/RNase-free water before quantitative PCR. RT–qPCR was then performed using the One-Step TB Green PrimeScript PLUS RT–PCR Kit (Takara Bio, Shiga, Japan) on a Thermal Cycler Dice Real-Time System (Takara Bio). The thermal profile consisted of reverse transcription at 42 °C for 5 min, an initial denaturation at 95 °C for 10 s, and 40 cycles of amplification at 95 °C for 5 s and 60 °C for 30 s. A melt-curve step was subsequently performed (95 °C for 15 s, 60 °C for 30 s, and 95 °C for 15 s). Relative mRNA expression was calculated using the ΔΔCt method and normalized to that of beta-actin as the internal control, using RNA from the control group as the calibrator sample.

#### 2.2.8. Short-Chain Fatty Acid Analysis

Short-chain fatty acid (SCFA) and organic acid analyses were outsourced to [excluded to facilitate double anonymization]. Acetic, propionic, succinic, lactic, n-butyric, and n-valeric acids were quantified in the fecal samples. The samples were shipped to the contractor under frozen conditions. Quantification was performed by [excluded for double anonymization] and measured by high-performance liquid chromatography using their validated analytical procedure. Concentrations were calculated using calibration with authentic standards and expressed as per sample mass.

#### 2.2.9. Gut Microbiota Analysis

Microbial DNA was extracted from the samples using a standard procedure at [excluded to facilitate double anonymization]. The bacterial *16S* rRNA gene was amplified and sequenced using high-throughput sequencing. Sequence data processing and downstream analyses were conducted using QIIME 2 [12], an open-source pipeline that integrates the steps required for amplicon sequence analysis from representative sequence inference to diversity analysis. In QIIME 2, representative sequences were inferred using an amplicon sequence variant-based approach, which statistically distinguishes true biological sequence variants from PCR/sequencing errors and provides higher taxonomic resolution than operational taxonomic unit clustering. Taxonomic assignments were performed using the SILVA reference database. Alpha-and beta-diversity metrics were calculated using QIIME 2. Alpha diversity was assessed using the Chao1, Shannon, and Simpson indices, while beta diversity was evaluated using Bray–Curtis and weighted UniFrac distances. Outputs were generated in the QIIME 2 visualization format (qzv) for an interactive review using QIIME 2 View [13,14].

#### 2.2.10. Plasma Biochemical Analysis

Blood plasma samples were analyzed by [excluded to facilitate double anonymization]. The total protein (TP), albumin (ALB), urea nitrogen (UN), creatinine (CRE), creatine kinase (CK), aspartate aminotransferase (AST), alanine aminotransferase (ALT), lactate dehydrogenase (LDH), and lactic acid (LA) levels were measured in each sample. Data are presented as the mean ± SEM for each group.

### 2.3. Ex Vivo Experiments (Peritoneal Macrophages)

#### 2.3.1. Isolation and Culture of Peritoneal Macrophages

Peritoneal macrophages were extracted from male C57BL/6J mice aged 7–8 weeks old. RPMI-1640 (Gibco) supplemented with 10% fetal bovine serum and penicillin–streptomycin (final concentration, 100 U/mL; FujiFilm Wako, Osaka, Japan) was injected into the peritoneal cavity (10 mL). After gently massaging the abdomen, the lavage fluid was collected and centrifuged at 350 × *g* for 5 min to obtain the peritoneal exudate cells. The cells were resuspended in the same RPMI-1640 complete medium, seeded in 24-well plates at a density of 1×10^5^ cells/well, and incubated overnight. Non-adherent cells were removed by washing, and adherent cells were used as peritoneal macrophages.

#### 2.3.2. Stimulation Protocol and RNA Collection

Macrophages were stimulated with heat-killed D279 LAB (10 µg/mL) for 24 h. Without removing the medium, LPS was added to a final concentration of 10 ng/mL, and the cells were incubated for 6 h. Cells were washed with PBS and harvested for RNA extraction using Quick-RNA MiniPrep Plus (Zymo Research) and subsequent qPCR analysis, as described in section 2.2.7.

### 2.4 In Vitro Experiments (IEC-6 Cells)

#### 2.4.1. IEC-6 Cell Culture

The IEC-6 rat intestinal epithelial cell line was obtained from the RIKEN Cell Bank (Tsukuba, Japan). The cells were cultured in Dulbecco’s modified Eagle’s medium (FujiFilm Wako) supplemented with 5% fetal bovine serum (Biowest, Nuaillé, France), 4 µg/mL insulin (FujiFilm Wako), and 1% penicillin–streptomycin (FujiFilm Wako) and incubated at 37 °C in a humidified atmosphere with 5% CO_2_.

#### 2.4.2. Proliferation Assay

IEC-6 cells were seeded in 96-well plates (8,000 cells/well) and cultured for 24 h before exposure. The cells were subsequently incubated with heat-killed D279 LAB for 48 h at specified concentrations. Cell proliferation was evaluated using the Cell Counting Kit-8 (CCK-8; Dojindo, Kumamoto, Japan) according to the manufacturer’s instructions. A microplate reader (Multimode Plate Reader ARVO X3; PerkinElmer, Waltham, MA, USA) was used to measure absorbance at 450 nm. Cell proliferation is expressed as a percentage of the untreated control.

#### 2.4.3. Evaluation of Barrier Function (TEER Assay)

IEC-6 cells were seeded onto Transwell inserts (Ad-MED Vitrigel 2, 12 well; Kanto Chemical, Tokyo, Japan) at a density of 1×10^5^ cells/insert and cultured for three days until a confluent monolayer was established, as confirmed by microscopy. The TEER was measured using a TEER measurement device (Kanto Chemical).

On Day 0, the baseline TEER values were measured before treatment. D279 was added to the Transwell inserts at concentrations of 0, 25, 50, and 100 μg/mL (n = 3). After 48 h of D279 treatment, the TEER was measured on day 3 to evaluate the effects of D279 on the epithelial barrier. After the day 3 TEER measurements, LPS was added to the LPS control and each D279-treated well at a final concentration of 100 µg/mL. After 48 h of incubation, TEER was measured on day 5 to evaluate the protective effects of D279 against LPS-induced epithelial barrier impairment.

#### 2.4.4. Quantitative PCR Analysis of IEC-6 Cells

After treatment, total RNA was isolated using the ISOSPIN Cell and Tissue RNA Kit (Nippon Gene, Tokyo, Japan) according to the manufacturer’s instructions. Gene expression was subsequently measured as described in section 2.2.7. The transcript levels of TJ-related genes were quantified using gene-specific primer sets. Expression values were normalized to those of *Gapdh*, and relative changes were calculated using the 2^−ΔΔCt^ method.

#### 2.4.5. Immunofluorescence Staining

After the specified treatments, the IEC-6 cells on the chamber slides were carefully washed with PBS. They were then fixed in 4% paraformaldehyde for 10 min at 25 °C. The cells were then rinsed thrice with PBS and treated with 0.1% Triton X-100 for 10 min at 25 °C to ensure permeability. The cells were incubated with 5% donkey serum for 1 h at 25 °C to prevent nonspecific binding. Cells were then incubated with Claudin 1 polyclonal antibody (Proteintech, Rosemont, IL, USA) at 1:200 dilution in blocking buffer for overnight at 4 °C. After three washes with PBS, samples were incubated with CoraLite594-conjugated ZO-1 polyclonal antibody (Proteintech) at 1:200 dilution in blocking buffer for 2 h, followed by CoraLite488 Donkey Anti-Rabbit IgG antibody (Proteintech) at 1:300 diluation for 1 h at 25 °C in the dark. After three additional washes with PBS, the nuclei were counterstained with DAPI for 5 min. The slides were maintained in PBS and subsequently imaged using a fluorescence microscope to ensure consistent acquisition settings across all experimental groups.

### 2.5. Statistical Analysis

Statistical analyses were conducted using SigmaPlot version 15.0 or Microsoft Excel. The Shapiro– Wilk test was applied to assess data normality, and the Brown–Forsythe test was used to evaluate equality of variances among groups. Statistical differences among multiple groups were determined using one-way analysis of variance (ANOVA), with subsequent pairwise comparisons performed using the Holm–Šídák method. Where appropriate, paired Student’s *t*-tests were employed to assess differences between matched observations. Significance was defined as *p* < 0.05.

## 3. Results

### 3.1. Effects of D279 on Body Weight Following LPS Challenge

Body weight was monitored throughout the study period to evaluate the effects of D279 supplementation and LPS challenge (Figure 1B). Body weight remained stable during the powdered diet transition and D279 supplementation period (days 0–14), with no significant differences among groups. After i.p. administration of LPS, body weight decreased markedly in both the LPS and LPS+D279 groups. On Day 15, body weight decreased by 8.0% and 3.4% from the pre-challenge baseline in the LPS and LPS+D279 groups, respectively (Figure 1C). However, no significant differences were observed between the LPS and LPS+D279 groups on days 16–18.

Daily food intake remained relatively stable before LPS challenge but decreased markedly after LPS administration (Table 3). On Day 18, food intake decreased to 50% and 54% of the control levels in the LPS and LPS+D279 groups, respectively.

**Table 1:**
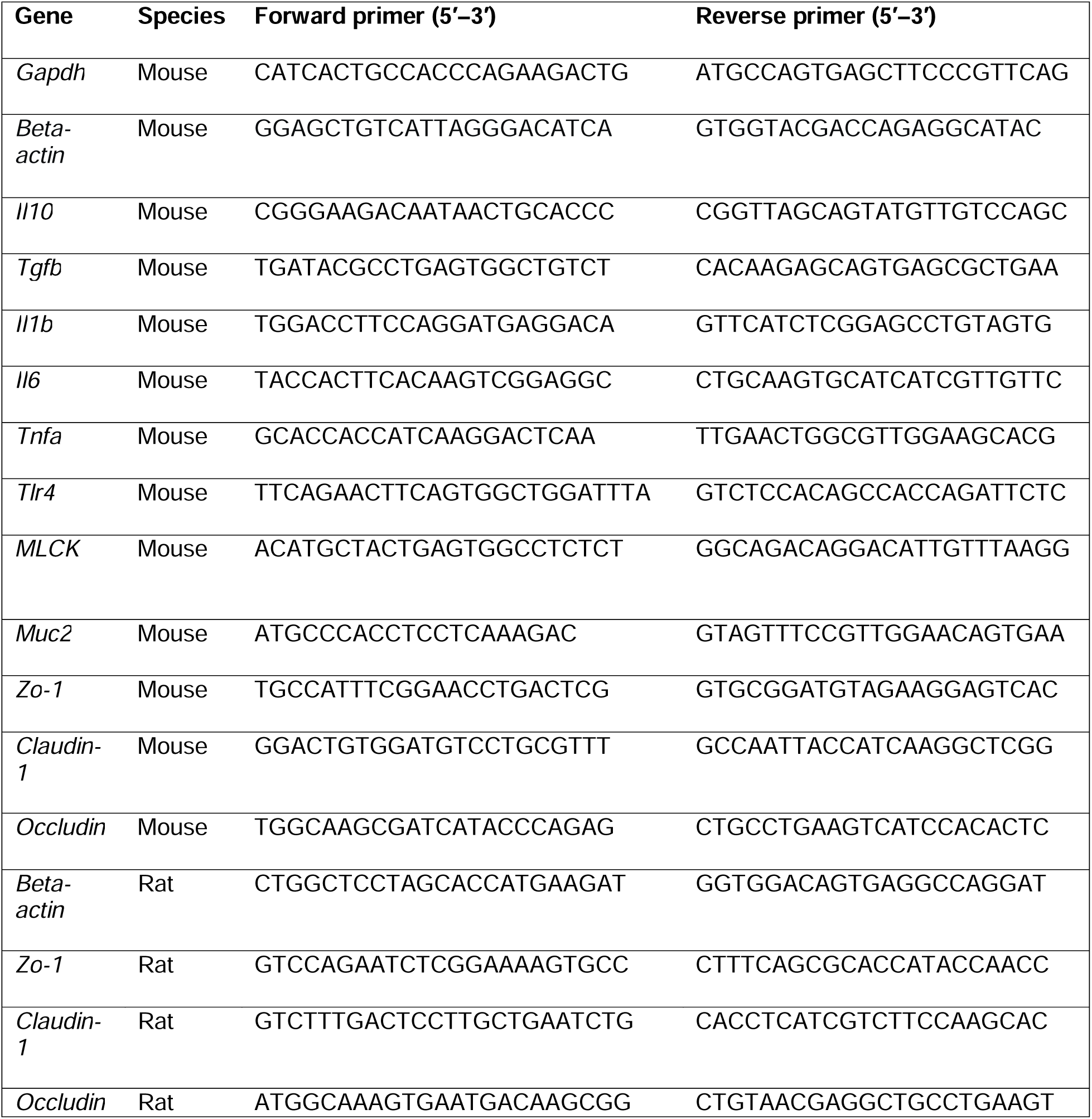
Forward and reverse primer sequences (5′–3′) used for RT–qPCR analysis, listed by target gene and species.

**Table 2:**
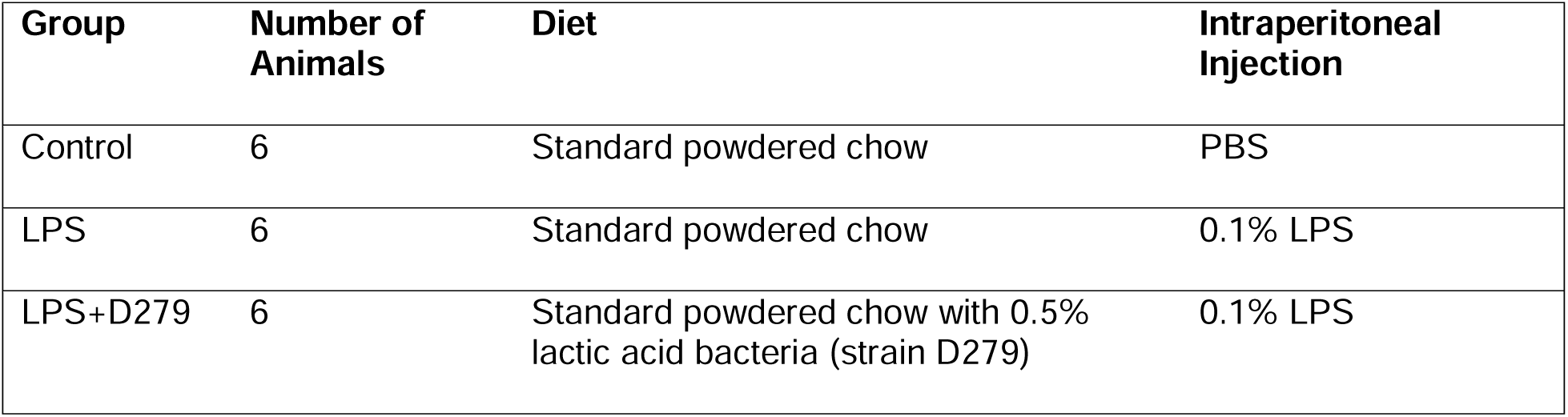
Grouping, diet, and intraperitoneal administration in animal experiments.

| **Group** | **Number of Animals** | **Diet** | **Intraperitoneal Injection** |
|---|---|---|---|
| Control | 6 | Standard powdered chow | PBS |
| LPS | 6 | Standard powdered chow | 0.1% LPS |
| LPS+D279 | 6 | Standard powdered chow with 0.5% lactic acid bacteria (strain D279) | 0.1% LPS |

**Table 3:** Daily food intake per mouse (g/mouse/day) in each group.

| **Group** | **Day 0** | **Day 7** | **Day 11** | **Day 14** | **Day 18** |
|---|---|---|---|---|---|
| **Control** | 3.10 ± 0.10 | 3.00 ± 0.10 | 3.00 ± 0.10 | 3.00 ± 0.10 | 2.80 ± 0.10 |
| **LPS** | 3.00 ± 0.06 | 3.10 ± 0.13 | 2.90 ± 0.14 | 2.90 ± 0.19 | 1.40 ± 0.12 |
| **LPS + D279** | 3.00 ± 0.00 | 3.00 ± 0.05 | 2.90 ± 0.08 | 2.90 ± 0.08 | 1.50 ± 0.41 |

Data are presented as the mean ± SD (cages per group).

**Table 4:** Plasma biochemical parameters in mice following LPS challenge and D279 supplementation.

|  | **TP (g/dL)** | **ALB (g/dL)** | **UN (mg/dL)** | **CRE (mg/dL)** | **AST (IU/L)** | **ALT (IU/L)** | **LDH (IFCC) (IU/L)** | **CK (IU/L)** | **LA (mg/dL)** |
|---|---|---|---|---|---|---|---|---|---|
| Control | 4.7 ± 0.1 | 3.0 ± 0.1 | 29.7 ± 3.8 | 0.11 ± 0.01 | 33.3 ± 2.8 | 17.3 ± 2.6 | 155.0 ± 21.2 | 49.0 ± 4.1 | 65.3 ± 11.2 |
| LPS | 4.4 ± 0.1 | 2.5 ± 0.1 | 24.6 ± 2.1 | 0.12 ± 0.01 | 58.3 ± 3.4 | 13.3 ± 0.8 | 237.0 ± 38.9 | 96.7 ± 4.4 | 63.8 ± 8.3 |
| LPS + D279 | 4.5 ± 0.5 | 2.5 ± 0.1 | 27.0 ± 3.5 | 0.13 ± 0.02 | 58.0 ± 9.2 | 16.6 ± 3.9 | 241.6 ± 48.0 | 96.0 ± 25.2 | 69.4 ± 10.0 |
| LPS vs Control (*p* value) | >0.001 | >0.001 | 0.008 | 0.036 | >0.001 | 0.002 | 0.001 | >0.001 | 0.396 |
| LPS+D279 vs LPS (*p* value) | 0.259 | 0.450 | 0.095 | 0.208 | 0.194 | 0.037 | 0.432 | 0.168 | 0.168 |
| LPS+D279 vs Control (*p* value) | 0.257 | >0.001 | 0.130 | 0.039 | >0.001 | 0.359 | 0.002 | 0.022 | 0.273 |

Data were obtained from six animals per group, except for the LPS+D279 group (n = 5), in which blood collection was unsuccessful in one animal. Values are presented as the mean ± SD. TP, total protein; ALB, albumin; UN, urea nitrogen; CRE, creatinine; AST, aspartate aminotransferase; ALT, alanine aminotransferase; LDH, lactate dehydrogenase; CK, creatine kinase; LA, lactate. Statistical comparisons were conducted between the LPS and Control, LPS+D279 and LPS, and LPS+D279 and Control groups.

### 3.2. LPS-Induced Colonic Inflammation

Macroscopic changes, histological alterations, and expression of colonic cytokine genes were assessed to evaluate the inflammatory and injurious effects of LPS on the colon. Representative images show an apparent reduction in colon length in LPS-challenged mice compared to that in the control group (Figure 2A). LPS significantly shortened the colon to 91% of the control value, whereas D279 supplementation attenuated this reduction, restoring colon length to 99% of the control value.

**Figure 2.**
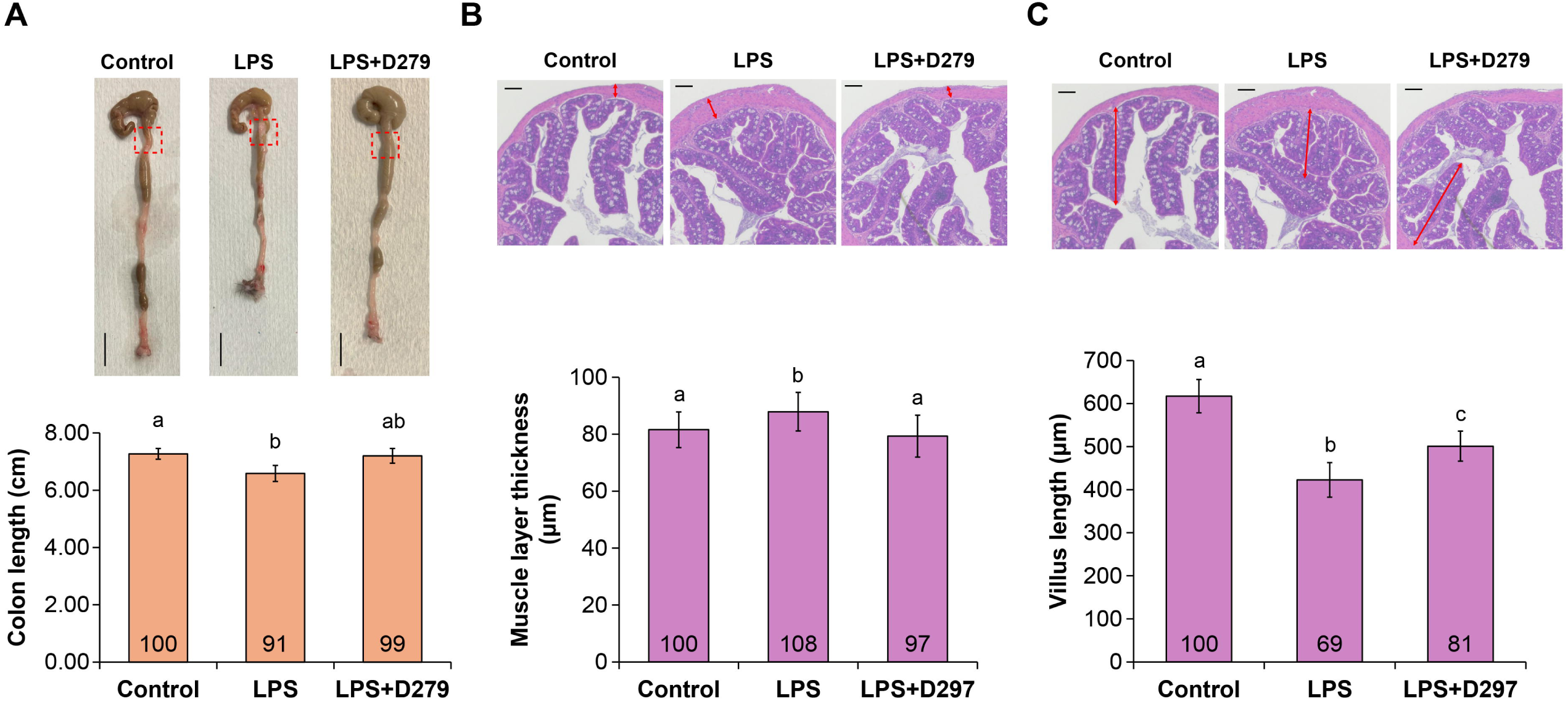
Heat-killed *Latilactobacillus sakei* D279 attenuates LPS-associated colon injury. (A) Representative images of the colons of mice in the Control, LPS, and LPS + D279 groups. Red boxes indicate the regions used for histological evaluation (scale bar = 1 cm). The bar graph shows the mean colon length (cm) measured from the cecum to the anus at necropsy. (B) Representative H&E-stained colon sections (scale bar = 0.1 mm). The bar graph shows the mean muscle layer thickness (μm). (C) Representative H&E-stained colon sections showing villus morphology (scale bar = 0.1 mm). The bar graph shows the mean villus length (μm). Data are presented as the mean ± SE; numbers within bars indicate values normalized to those of the control group (control = 100%). Bars with different letters indicate significant differences among groups (one-way ANOVA followed by pairwise multiple comparisons using the Holm–Sidak method; *p* < 0.05). ANOVA, analysis of variance; H&E, hematoxylin and eosin; LPS, lipopolysaccharide

Histological evaluation further indicated structural changes in the colon after LPS challenge. Specifically, LPS significantly increased muscle layer thickness to 108% of the control value, whereas D279 supplementation significantly reversed this effect (Figure 2B). In addition, LPS challenge significantly reduced the villus length to 69% of the control value, whereas D279 supplementation partially attenuated this reduction, with the villus length reaching 81% of the control value (Figure 2C).

RT–qPCR analysis indicated that LPS caused a notable increase in both pro-inflammatory (*Tnfa*, *Il6*, *Il1b*, and *Il2b*) and anti-inflammatory (*Il10* and *Tgfb*) cytokine mRNA expression in the colon compared to that in the control group (Figure 3). This was consistent with the macroscopic and histological results. The LPS-induced increase in cytokine transcript levels was attenuated in the D279 supplementation group. Collectively, these findings indicate that LPS challenge induces colonic inflammatory and structural alterations characterized by colon shortening, muscle layer thickening, villus shortening, and increased cytokine gene expression, whereas D279 supplementation attenuates these alterations.

**Figure 3.**
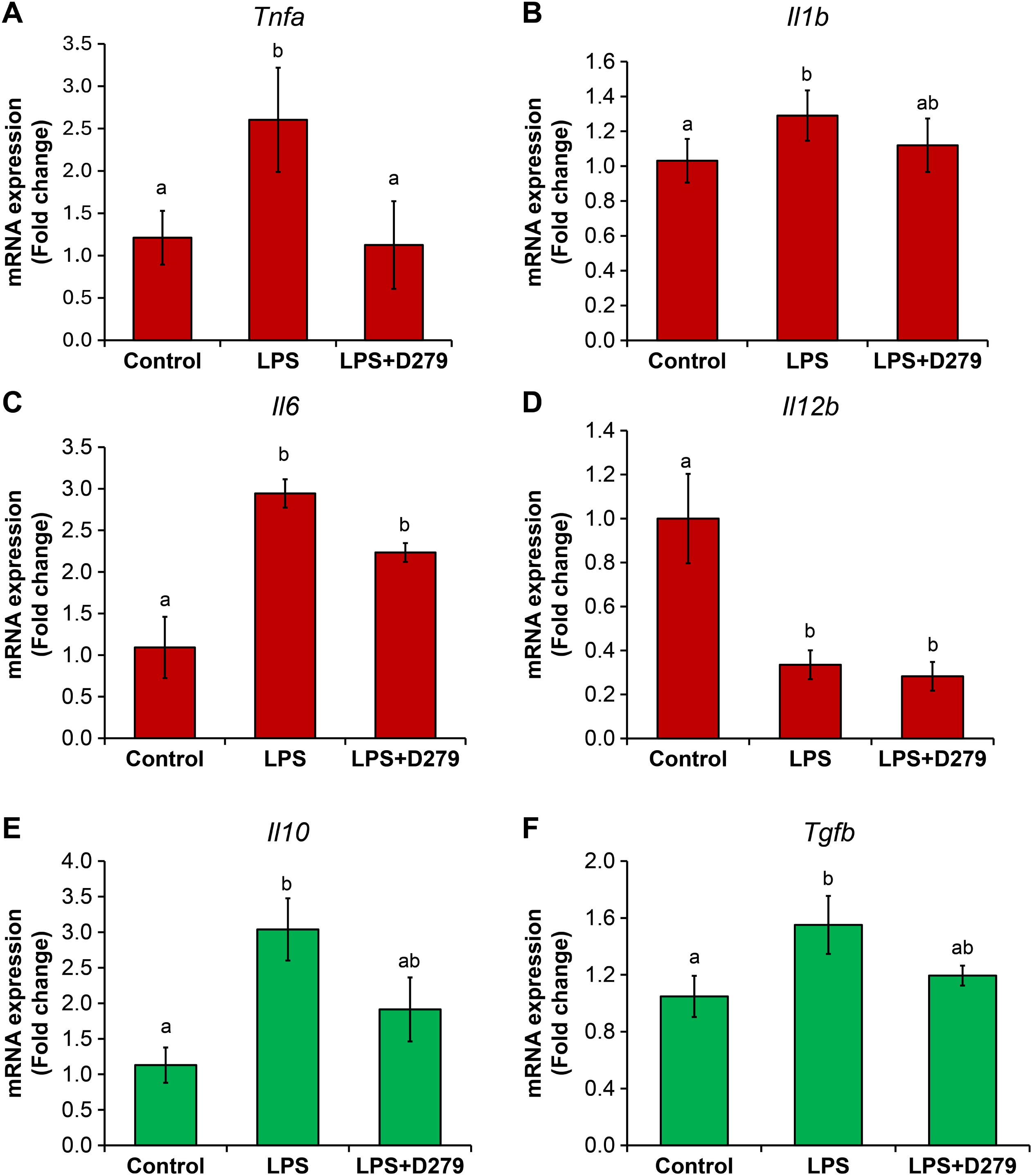
Effects of D279 supplementation on the expression of inflammatory cytokines in the colon of mice. The relative mRNA expression levels of pro-inflammatory cytokines, including (A) *Tnfa*, (B) *Il6*, (C) *Il1b*, and (D) *Il12b*, as well as anti-inflammatory cytokines, such as (E) *Il10* and (F) *Tgfb*, were assessed in colonic tissue. Data are expressed as the mean ± SE. Bars marked with different letters denote significant differences between groups, as determined by one-way ANOVA followed by pairwise multiple comparisons using the Holm–Sidak method, with a significance level of *p* < 0.05. ANOVA, analysis of variance

### 3.3 D279 Attenuates LPS-associated Disruption of Intestinal Barrier Integrity

Goblet cell abundance was assessed using AB-PAS staining to further evaluate the effects of D279 on the intestinal barrier integrity (Figure 4A). LPS administration significantly reduced the abundance of goblet cells in the intestinal mucosa compared to that in the control group, indicating LPS-induced goblet cell depletion. D279 supplementation attenuated this reduction, and goblet cell abundance remained higher than that observed in the LPS group.

**Figure 4.**
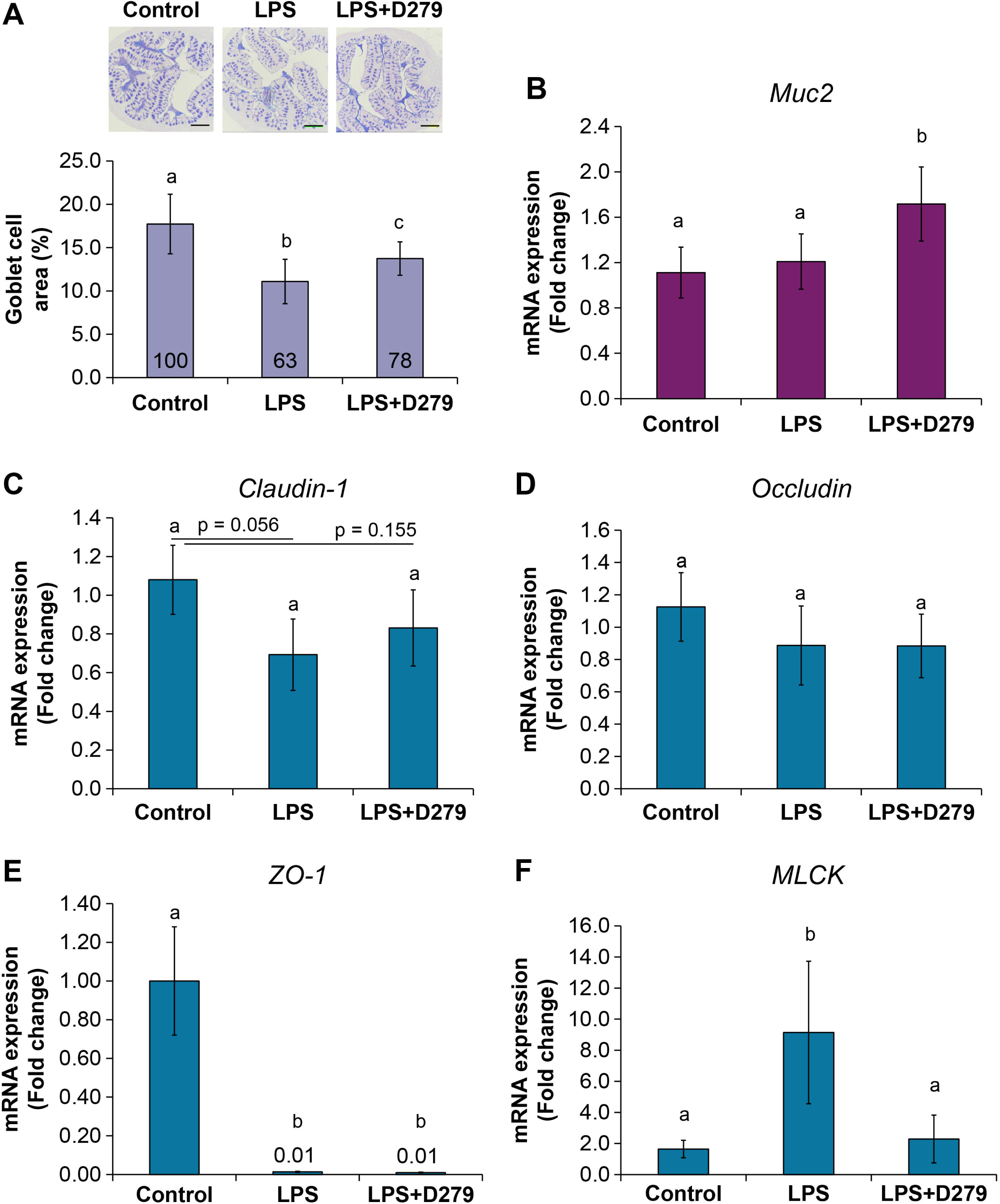
D279 preserves colonic mucosal integrity and modulates barrier-related markers after systemic LPS challenge. (A) Representative histological images of distal colon sections (stained to visualize the mucosal architecture and goblet cells) of the control, LPS, and LPS+D279 groups (top). The bar graphs show quantification of barrier-and inflammation-associated readouts. Data are presented as the mean ± SE. Scale bar indicates the micrograph. LPS, lipopolysaccharide

Next, we examined the expression of MUC2, a major mucin produced by the intestinal goblet cells (Figure 4B). LPS administration did not significantly alter *Muc2* mRNA expression compared with that in the control group. However, D279 supplementation in LPS-treated mice significantly increased *Muc2* expression compared to that in the LPS group. Despite the reduction in goblet cell abundance following LPS exposure, LPS alone did not significantly affect *Muc2* transcription, whereas D279 promoted *Muc2* expression under inflammatory conditions.

The expression of genes related to epithelial TJ and barrier regulation was examined using qPCR. Although LPS administration reduced the mRNA expression of *Claudin-1* and *Occludin*, these changes were not significant (Figure 4C and 4D). *ZO-1* expression was significantly reduced by LPS treatment, whereas D279 supplementation did not significantly restore *ZO-1* expression (Figure 4E). However, *MLCK* expression was significantly increased following LPS administration, and D279 treatment significantly attenuated this increase (Figure 4F).

Collectively, these findings indicate that LPS-induced intestinal barrier disruption is accompanied by goblet cell depletion, reduced *ZO-1* expression, and increased *MLCK* expression. D279 partially preserved goblet cell abundance, enhanced *Muc2* expression, and suppressed the LPS-induced increase in *MLCK*, suggesting that this strain supports both mucus and epithelial barrier integrity during LPS-induced intestinal inflammation.

### 3.4. D279 Supplementation Alters the Gut Microbial Community Structure

The composition and diversity of the gut microbiota were analyzed to determine whether D279 supplementation affects the intestinal microbial community under LPS-induced inflammatory conditions. The relative abundances of the predominant bacterial taxa differed among the control, LPS, and LPS+D279 groups (Figure 5A). Barnesiellaceae (36.4%), Lactobacillaceae (17.0%), and Lachnospiraceae (14.9%) were the predominant families in the control group. Following LPS administration, the relative abundance of Barnesiellaceae and Lactobacillaceae decreased to 27.7% and 10.1%, respectively. In contrast, Clostridia UCG-014 levels increased from 2.6% in the control group to 11.0% in the LPS group. The relative abundance of Lactobacillaceae in the LPS+D279 group markedly increased to 34.1%, whereas that of Clostridia UCG-014 decreased to 5.9%. The abundance of Barnesiellaceae further decreased to 20.0% in the LPS+D279 group. These findings indicate that LPS and D279 supplementation were associated with a distinct shift in the relative composition of the gut microbiota.

**Figure 5.**
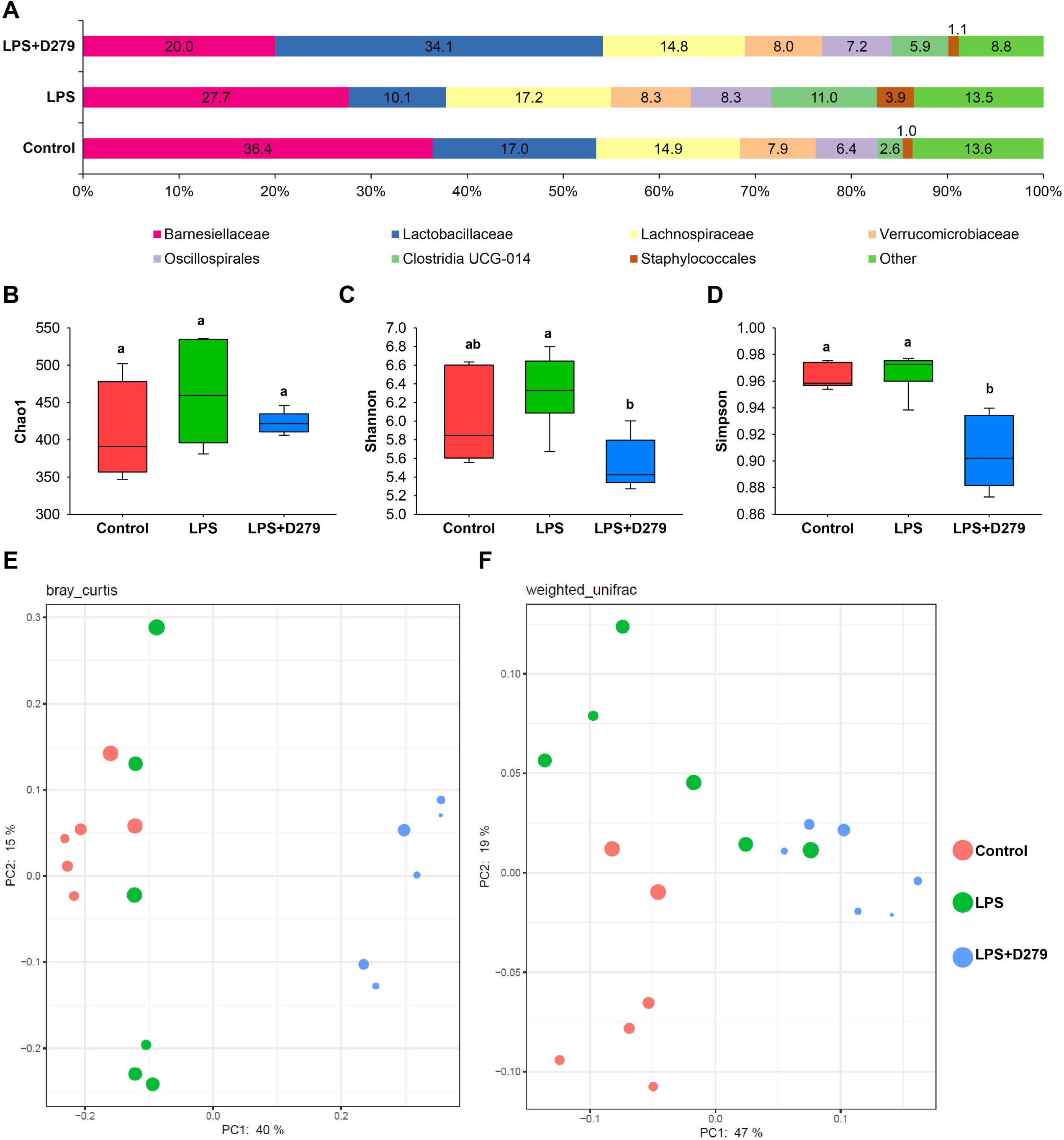
**D279 supplementation alters gut microbial community structure following systemic LPS challenge**. (A) Relative abundance of the predominant bacterial taxa in fecal samples from the Control, LPS, and LPS+D279 groups. (B–D) Alpha diversity, as assessed using the Chao1, Shannon, and Simpson indices, respectively. Statistical differences in alpha diversity were evaluated using the Kruskal–Wallis test. (E) PCoA based on Bray–Curtis dissimilarity and (F) weighted UniFrac distances, showing differences in overall gut microbial community structure among the three groups. Differences in beta diversity among groups were assessed using analysis of similarities. Each point represents an individual mouse. LPS, lipopolysaccharide; PCoA, principal coordinate analysis

Alpha diversity was assessed using the Chao1, Shannon, and Simpson indices (Figure 5B-D). The Chao1 index revealed no significant differences between the three groups, suggesting that the estimated microbial richness was not notably affected by LPS administration or D279 supplementation. Conversely, the Shannon index was significantly reduced in the LPS+D279 group compared to that in the LPS group. Similarly, the Simpson index was significantly lower in the LPS+D279 group than that in the Control and LPS groups. Therefore, although the estimated microbial richness remained constant, D279 supplementation was linked to modifications in the distribution and relative abundance of bacterial taxa, leading to changes in the overall alpha diversity.

Beta diversity was evaluated using Bray–Curtis and weighted UniFrac distances, as depicted in Figure 5E,F. Principal coordinate analysis demonstrated a distinct separation between the control, LPS, and LPS+D279 groups for both distance measures. For Bray–Curtis distances, pairwise analysis of similarities demonstrated significant differences between the Control and LPS (R = 0.648, q = 0.004), Control and LPS+D279 (R = 1.000, q = 0.004), and LPS and LPS+D279 (R = 0.883, q = 0.004) groups. Similarly, weighted UniFrac analysis showed significant differences between the Control and LPS (R = 0.528, q = 0.003), Control and LPS+D279 (R = 0.993, q = 0.004), and LPS and LPS+D279 (R = 0.628, q = 0.003) groups. These results demonstrate that LPS administration altered the overall gut microbial community structure, and that D279 supplementation further shifted the community to a composition distinct from that of both the Control and LPS groups.

Fecal SCFA levels were analyzed to determine whether these microbial alterations were accompanied by changes in the secretion of microbial metabolites (Figure 6). LPS administration did not significantly affect acetic, propionic, or succinic acid levels. In contrast, lactic, n-valeric, and n-butyric acid levels were significantly reduced to approximately 23%, 61%, and 58% of the control levels, respectively. D279 supplementation partially attenuated the LPS-associated reduction in n-valeric and n-butyric acids, with levels reaching approximately 73% and 72% of the control levels, respectively. However, LA levels were not similarly preserved by D279 supplementation. Together, these findings demonstrate that LPS-induced inflammation was accompanied by alterations in the gut microbial composition and selected fecal SCFAs, whereas D279 supplementation induced a distinct microbial community structure and partially preserved specific SCFAs.

**Figure 6.**
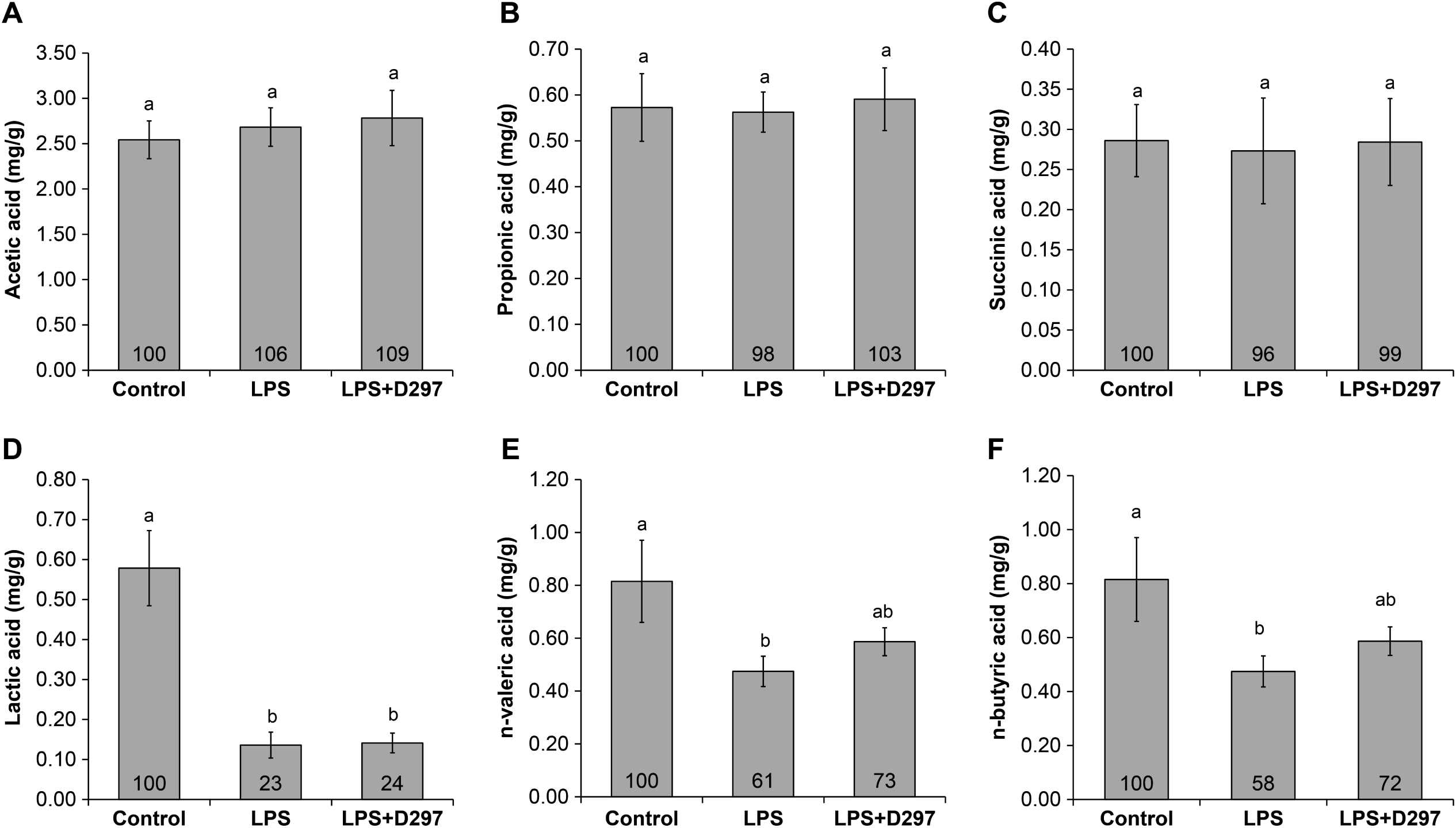
Effects of heat-killed *L. sakei* D279 on fecal SCFA levels in mouse experiments. (A– F) Fecal concentrations of individual SCFAs (acetic acid, propionic acid, succinic acid, lactic acid, n-valeric acid, and n-butyric acid) were measured and are expressed as the mean (mg/g feces) for each group. Numbers above the bars indicate the relative SCFA levels compared to those in the control group, which was set at 100%. Data are presented as the mean ± SE (n = 6 or 5 per group). SCFA, short-chain fatty acid

### 3.5 LPS-Associated Plasma Biochemical Alterations

Plasma biochemical parameters were analyzed to assess the systemic responses to LPS administration and the effects of D279 supplementation. LPS administration significantly decreased TP (*p* < 0.001) and ALB (*p* < 0.001) levels and significantly increased AST, LDH, and CK activities (*p* < 0.001, *p* = 0.001, and *p* < 0.001, respectively) compared to those in the control group. UN and CRE levels were also significantly altered by LPS administration (*p* = 0.008 and *p* = 0.036, respectively), whereas plasma LA levels did not significantly differ between the LPS and control groups (*p* = 0.396).

D279 supplementation did not significantly attenuate LPS-induced changes in ALB, CRE, AST, LDH, and CK levels (*p* > 0.05 for all comparisons with the LPS group). TP and UN levels showed a numerical recovery toward the control level in the LPS + D279 group (4.5 and 27.0 mg/dL) compared with that in the LPS group (4.4 and 24.6 mg/dL), although the difference was not statistically significant (*p* = 0.095). Furthermore, ALT activity was significantly higher in the LPS + D279 group than in the LPS group (16.6 vs. 13.3 IU/L, *p* = 0.037), and the levels were comparable to those of the control group (17.3 IU/L; *p* = 0.359). Plasma LA levels remained unchanged following D279 supplementation (*p* = 0.168 vs. LPS; *p* = 0.273 vs. the control group).

Overall, D279 showed limited effects on most LPS-associated systemic biochemical alterations but significantly attenuated the LPS-induced decrease in ALT activity, resulting in levels comparable to those of the control group.

### 3.6. D279 Pretreatment Modulates LPS-Induced Inflammatory Responses in Ex Vivo Mouse Peritoneal Macrophages

To assess whether D279 modulates macrophage inflammatory responses, mouse peritoneal macrophages were pretreated with D279 for 24 h and subsequently stimulated with LPS for 6 h. Cytokine and LPS receptor gene expression was evaluated using RT–qPCR (Figure 7).

**Figure 7.**
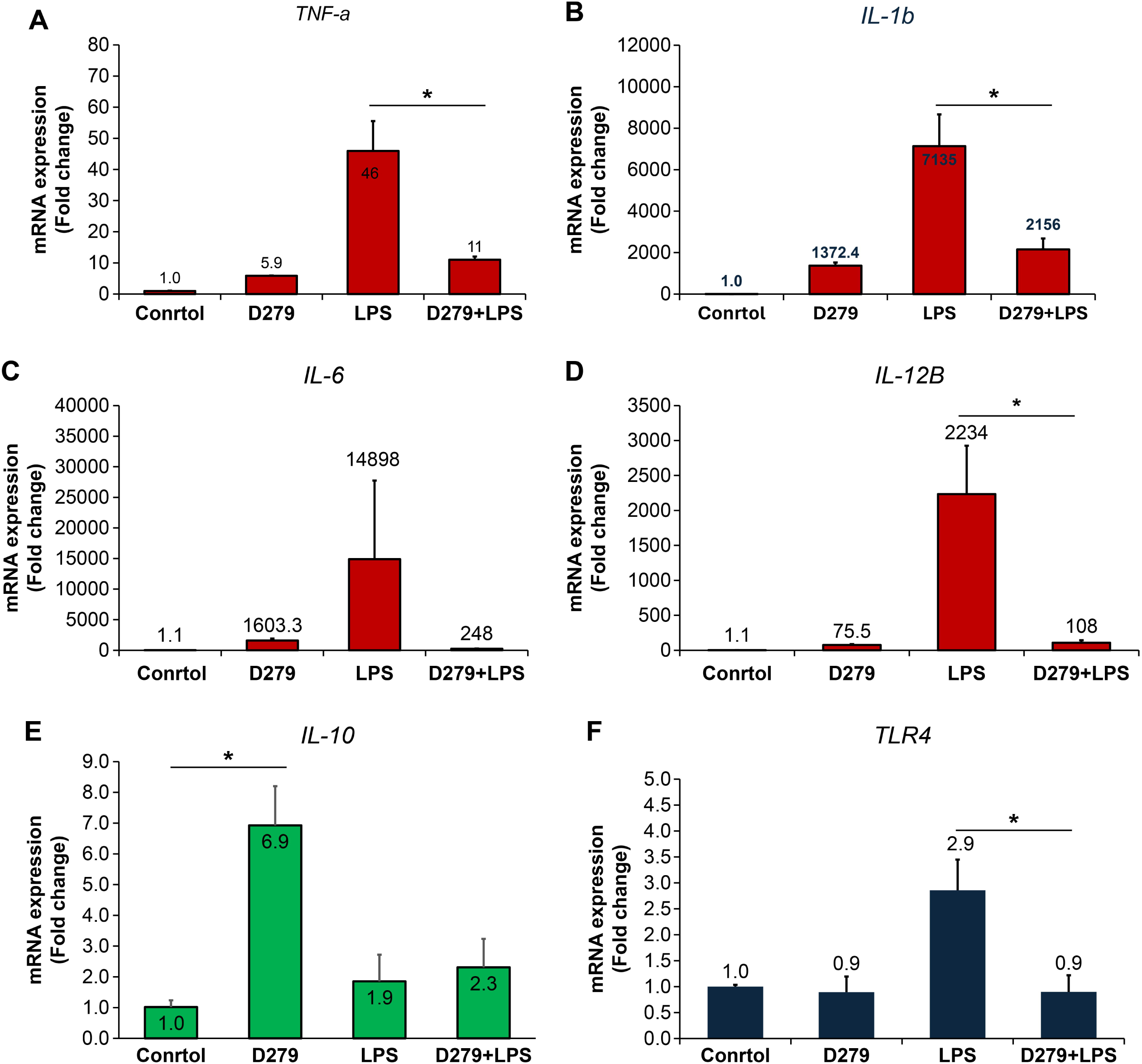
Effects of heat-killed *L. sakei* D279 on inflammatory gene expression in ex vivo macrophages. The mRNA expression levels of inflammation-related genes (A–E: *Tnfa*, *Il1b*, *Il6*, *Il12b*, *Il10*) and (F) the LPS signaling receptor *Tlr4*, as analyzed by quantitative real-time PCR and normalized to the control group. LPS, lipopolysaccharide

D279 treatment alone increased the expression of pro-inflammatory cytokines compared to that in untreated macrophages, with *Tnfa*, *Il1b*, *Il6*, and *Il12b* reaching approximately 6-, 1,372-, 1,603-, and 76-fold the values of the untreated control, respectively. LPS stimulation induced a substantially greater inflammatory response, increasing the expression of *Tnfa*, *Il1b*, *Il6*, and *Il12b* to approximately 46-, 7,135-, 14,898-, and 2,234-fold, respectively. Moreover, D279 pretreatment markedly attenuated this LPS-induced response, with the expression levels in the LPS + D279 group reaching approximately 11-, 2,156-, 248-, and 108-fold of the untreated control for *Tnfa*, *Il1b*, *Il6*, and *Il12b*, respectively. Thus, although D279 alone stimulated cytokine expression under basal conditions, D279 pretreatment substantially reduced the magnitude of the inflammatory response elicited by subsequent LPS stimulation.

D279 treatment alone also increased *Il10* expression by approximately 7-fold compared to that in the untreated control. In contrast, *Il10* expression in both the LPS and LPS + D279 groups remained at an approximately 2-fold increase compared to that in the untreated control. This indicates that the D279-induced increase in *Il10* under basal conditions was not maintained following LPS stimulation. Furthermore, LPS increased *Tlr4* expression by approximately 3-fold compared to that in the untreated control, whereas *Tlr4* expression in the D279-alone and LPS + D279 groups remained comparable to that in the untreated control.

Collectively, these findings indicate that D279 exerts context-dependent immunomodulatory effects on macrophages. D279 alone activated cytokine-related gene expression and increased *Il10* expression, whereas pretreatment with D279 markedly limited the excessive pro-inflammatory response induced by LPS and prevented LPS-associated upregulation of *Tlr4* expression.

### 3.7. D279 Improves the Proliferation of IEC-6 Cells

To investigate whether D279 protects intestinal epithelial cells against LPS-induced cellular stress, IEC-6 cells were pretreated with this strain for 24 h before LPS stimulation; cell viability was assessed using the CCK-8 assay. In the absence of LPS, D279 significantly increased the viability of IEC-6 cells in a concentration-dependent manner, reaching approximately 120–130% that of the untreated controls (Figure 8A). This effect was consistently observed in independent experiments.

**Figure 8.**
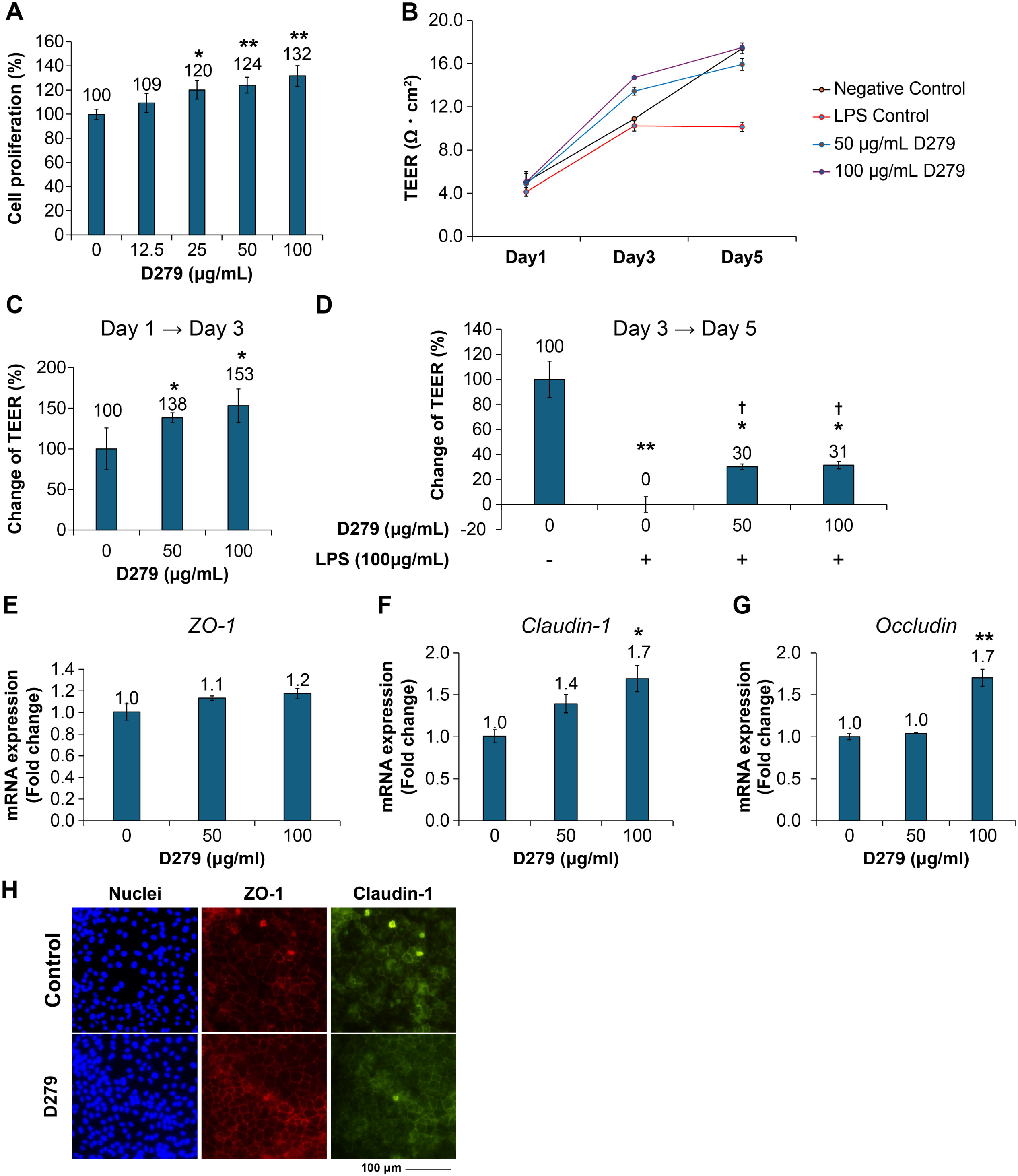
Effects of heat-killed *L. sakei* D279 on intestinal epithelial cell proliferation, barrier function, and TJ gene expression. (A) Proliferation of IEC-6 cells following treatment with heat-killed *L. sakei* D279 (0, 50, and 100 μg/mL). (B) TEER (Ω·cm²) was measured on days 1 (baseline, immediately before D279 treatment), 3 (immediately before LPS stimulation), and 5 (end of the experiment). TEER measurements were used to evaluate changes in epithelial barrier integrity following D279 treatment and subsequent LPS stimulation. (C, D) Percentage change in TEER following D279 treatment (day 1 to day 3; C) and subsequent LPS (100 μg/mL) challenge (day 3 to day 5; D), respectively. (E–G) mRNA expression of *Zo-1*, *Claudin-1*, and *Occludin i*n IEC-6 cells following D279 (100 μg/mL) treatment. Representative fluorescence images show the expression of the indicated markers. Data are presented as the mean ± SE. Statistical significance is indicated by *p* < 0.05 and \**p* < 0.01 compared with the untreated control (0 μg/mg D279), and by +*p* < 0.05 compared with the LPS-treated control (0 μg/mg D279 + LPS). (H) Representative immunofluorescence staining images of IEC-6 cells showing TJ protein expression following D279 treatment. LPS, lipopolysaccharide; TEER, transepithelial electrical resistance; TJ, tight junction

### 3.8. D279 Promotes Intestinal Epithelial Barrier Development and Attenuates LPS-Induced TEER Reduction

To investigate the effects of D279 on intestinal epithelial cells, IEC-6 cell proliferation was evaluated after 48 h treatment with increasing concentrations of D279. D279 increased cell proliferation in a concentration-dependent manner, indicating a positive effect on IEC-6 cell growth under the experimental conditions.

TEER was monitored during barrier formation and after LPS challenge to further determine whether D279 enhanced intestinal epithelial barrier function (Figure 8B). IEC-6 cells were treated with D279, and TEER was measured on days 1 and 3. Changes in TEER from Day 1 to day 3 were compared among the treatment groups. Compared with the untreated control, D279 significantly increased the change in TEER at 50 and 100 μg/mL, reaching approximately 138% and 153%, respectively (Figure 8C). These findings indicate that D279 promotes the development of epithelial barrier function during IEC-6 monolayer formation.

To examine the protective effect of D279 under inflammatory conditions, cells were pretreated with this strain before LPS stimulation on day 3, and TEER was measured on day 5. In the untreated control group, the change in TEER from day 3 to day 5 was defined as 100%. LPS markedly suppressed the increase in TEER, resulting in an approximately 0% change compared to that on day 3. In contrast, D279 pretreatment maintained the increase in TEER after LPS exposure, with TEER changes reaching approximately 30% and 31% at 50 and 100 μg/mL D279, respectively (Figure 8D). These findings indicate that D279 pretreatment attenuates LPS-induced impairment of epithelial barrier development.

Overall, D279 promoted IEC-6 cell proliferation and enhanced epithelial barrier development under basal conditions, whereas pretreatment with D279 partially preserved the TEER following LPS exposure.

### 3.9. D279 Enhances TJ-Related Gene Expression and Protein Localization

To further investigate the effects of D279 on intestinal epithelial barrier function, the expression and localization of TJ components were evaluated in IEC-6 cells treated with D279 (100 μg/mL). D279 treatment increased *ZO-1* mRNA expression by approximately 1.2-fold compared to that in the untreated control, although this increase was not significant (Figure 8E). In contrast, *Claudin-1* and *Occludin* mRNA expression significantly increased by approximately 1.7-fold following D279 treatment (Figure 8F and 8G).

Immunofluorescence analysis was performed on day 3, prior to LPS stimulation, using cells cultured in parallel with those used for the TEER experiment. The results revealed more pronounced membrane localization of Zo1 and Claudin-1 in D279-treated cells than in untreated cells (Figure 8H). Although TJ structures were still developing at this stage, the D279-treated cells exhibited clear membrane-associated localization of these proteins. Together, these findings indicate that D279 promotes the development of the intestinal epithelial barrier, accompanied by increased expression of key TJ-related genes and enhanced localization of TJ proteins to the cell membrane.

## Discussion

In the present study, oral administration of heat-killed *L. sakei* D279 attenuated LPS-induced colonic injury. Intraperitoneal LPS administration elicited marked inflammatory responses characterized by colon shortening and disruption of the colonic architecture, whereas D279 supplementation mitigated these pathological changes. D279 also partially prevented the LPS-induced reduction in colonic goblet cell abundance and increased the expression of the mucus-related gene *Muc2* in the colon. Furthermore, D279 supplementation modulated the colonic microbiota composition in LPS-challenged mice, suggesting the potential contribution of microbiota alterations to its protective effects. Consistent with these in vivo observations, experiments demonstrated that D279 enhanced the epithelial barrier function and attenuated LPS-induced TEER in vitro. In the ex vivo experiments, D279 modulated the expression of inflammation-related genes in macrophages, indicating a homeostatic priming response. Additionally, LPS stimulation markedly increased the expression of pro-inflammatory cytokines in macrophages, whereas D279 pretreatment attenuated this response, suggesting that LPS directly affects macrophage inflammatory responses. Collectively, these results suggest that D279 prevents systemic inflammation-associated colonic injury, potentially by preserving the gut barrier and mucus-associated defenses.

### 4.1. Study Overview and Relevance of the Systemic LPS Model

Intraperitoneal LPS administration primarily activates innate immune signaling via TLR4. It induces a systemic inflammatory response that can secondarily disrupt intestinal epithelial homeostasis [4,15,16]. In the present study, LPS induced measurable systemic physiological responses, as reflected by changes in plasma biochemical indices, while simultaneously inducing distal colonic injury. D279 did not substantially normalize most of these systemic biochemical alterations, suggesting that its most prominent protective effects were expressed at the intestinal/colonic level rather than through the broad suppression of systemic responses.

This model differs conceptually from that of dextran sulfate sodium-induced colitis, in which the initiating injury was primarily luminal or epithelial [17–18]. Therefore, the LPS model provides a complementary framework for evaluating whether an orally administered postbiotic can preserve intestinal integrity under systemic inflammatory stress, rather than only in models in which the intestinal mucosa is directly exposed to injurious stimuli.

### 4.2. Preservation of Colonic Structure and Mucosal Compartments

Systemic LPS challenge induces characteristic colonic pathology, including colon shortening and architectural disruption, together with impairment of mucosal protective compartments. These findings are consistent with those of previous studies in mice, showing that systemic LPS administration induces colonic mucosal damage, including villus disruption and goblet cell depletion [19–21]. D279 significantly preserved colon length and attenuated histological abnormalities, indicating that non-viable bacterial preparations can affect distal colonic integrity, despite systemic inflammatory triggers.

LPS also reduced the goblet cell count, whereas D279 treatment mitigated this reduction. In parallel, D279 increased colonic *Muc2* expression compared with treatment with LPS alone. Taken together, these findings suggested that D279 supports the preservation of mucus-associated defense mechanisms during inflammatory stress. However, as mucus layer thickness, mucus secretion, and mucin organization were not directly quantified, the present findings should be interpreted as evidence for the preservation of mucus-associated cellular and molecular components rather than definitive restoration of the physical mucus layer.

### 4.3. Epithelial Barrier Support and TJ regulation by D279: Integrating In Vitro and In Vivo Evidence

TJs are key components of the intestinal epithelial barrier. They regulate paracellular permeability and maintain an intact mucosal lining. Proteins linked to TJs, such as occludins, claudins, and zonula occludens (ZO), are crucial for the formation, organization, and maintenance of TJ complexes [22]. LPS-induced inflammatory responses disrupt TJ organization and impair epithelial barrier function, resulting in increased intestinal permeability. This facilitates the translocation of luminal microbial components, which may further exacerbate intestinal inflammation [23]. In this study, IEC-6 experiments provided mechanistic validation of barrier protection. D279 promoted TEER development under basal conditions and attenuated LPS-induced decline in TEER. D279 also increased IEC-6 proliferation, suggesting its potential role in epithelial maintenance. At the transcriptional level, D279 upregulated the expression of TJ-related mRNAs in IEC-6 cells, including *claudin*-1 and *occludin*. This is consistent with a pro-barrier epithelial response in vitro. Long-term D279 supplementation in healthy mice was also associated with increased expression of intestinal barrier-related genes in the small intestine (Supplementary Fig. S4). Although these findings do not directly demonstrate improved barrier function, they provide additional evidence that D279 affects epithelial barrier-associated pathways under basal physiological conditions.

The effects of D279 on the expression of TJ-related genes were less uniform in vivo. At the dose used in the present study, LPS significantly reduced colonic *Zo-1* expression, whereas *claudin-1* and occludin expression was not significantly altered. Although reductions in the expression of these TJ-related genes have been reported in other LPS models [24], differences in the LPS dose, administration route, and sampling time may have contributed to the distinct responses observed in this study. D279 did not significantly restore *Zo1* expression. However, it attenuated LPS-induced upregulation of *MLCK* expression, suggesting that its barrier-protective effects involve the regulation of epithelial permeability pathways rather than the broad restoration of TJ-related transcription. MLCK is a Ca²⁺/calmodulin-dependent protein kinase that controls intestinal epithelial permeability through phosphorylation of the myosin light chain (MLC), which induces perijunctional actomyosin ring contraction and TJ opening [25,26]. Increased *MLCK* expression and activity following LPS stimulation promotes MLC phosphorylation, leading to actomyosin contraction and cytoskeletal remodeling. These changes can alter TJ organization and increase paracellular permeability, thereby compromising the integrity of the intestinal barrier [27]. In this context, D279-mediated attenuation of LPS-induced upregulation of *MLCK* expression suggests a potential mechanism that contributes to preservation of the epithelial barrier. However, MLC phosphorylation and TJ protein localization were not directly assessed in the present study. Therefore, further studies are required to confirm whether D279 modulates this pathway at a functional level.

The differences between the in vitro and in vivo findings may reflect the greater complexity of the intestinal environment, in which barrier integrity is determined not only by epithelial transcription, but also by TJ protein localization, cytoskeletal regulation, mucus-associated protection, immune cell signals, and microbial interactions.

Taken together, the TEER measurements, TJ gene expression, and *MLCK* expression findings suggest that D279 supports intestinal barrier integrity through multiple complementary mechanisms rather than through the complete restoration of all TJ-related transcripts in vivo. The increased expression of barrier-associated genes observed in healthy mice further suggests that D279 affects epithelial barrier homeostasis independent of an overt inflammatory challenge. These findings demonstrate that the epithelial barrier function must be considered within the broader context of the coordinated epithelial, immune, mucus-associated, and microbial responses observed in the present study.

### 4.4. Context-Dependent Immunomodulation and Innate Immune Recognition of D279

The intestinal-protective phenotype was accompanied by immunomodulatory activity that is consistent with context-dependent regulation rather than constitutive anti-inflammatory signaling. D279 increased IL-10 expression under basal conditions and attenuated LPS-induced proinflammatory responses ex vivo. However, it did not increase IL-10 expression upon LPS stimulation. These findings suggest that D279 modulates the responsiveness of innate immune cells rather than acting solely through sustained induction of IL-10 [28,29].

Bacterial-derived bioactive factors activate TLR2 and induce macrophage cytokine responses in a species-and strain-dependent manner. This indicates the potential contribution of bacterial components to immunomodulation [30]. FITC-labeled D279 showed clear cell-associated fluorescence in macrophage-like cells, confirming a direct interaction between the bacterial preparation and macrophages (Supplementary Fig. S1). Furthermore, inhibition of TLR2 or Dectin-1 markedly reduced D279-induced IL-12p70 production, whereas that of TLR4 or TLR9 had a more limited effect (Supplementary Fig. S2). These findings suggest that TLR2-and Dectin-1-dependent recognition pathways contribute to macrophage responses to D279 rather than to classical LPS-associated TLR4 signaling. This interpretation is consistent with the established synergistic signaling between C-type lectin receptors and Toll-like receptors in innate immune recognition [31]. D279 also contained a relatively high level of lipoteichoic acid (LTA) compared with that in other bacterial strains. Its LTA content was comparable to that of the *L. sakei* type strain (Supplementary Fig. S3). Given the established role of LTA in bacterial recognition and immune activation, the relatively high LTA content in D279 may be related to its TLR2-associated recognition. However, the present study did not establish LTA as the definitive active ligand; other cell wall-associated components may have also contributed to the observed response.

Importantly, the ex vivo macrophage findings provide a mechanistic link between D279 exposure and the modulation of innate immune responses, whereas the in vivo results indicate that this immunomodulatory activity occurs in parallel with the preservation of colonic structure and barrier-associated functions. One plausible interpretation is that D279 modulates the inflammatory environment to which the intestinal epithelium is exposed during systemic inflammatory stress, indirectly contributing to epithelial and mucosal protection. This interpretation is consistent with the role of innate immune signaling in the regulation of inflammatory responses and tissue homeostasis [32].

### 4.5. Microbiota and SCFAs: Links to Intestinal Protection

In addition to the epithelial and immune effects, D279 supplementation was associated with changes in the intestinal microbiota. LPS altered the overall microbial community structure, and the LPS+D279 group showed a distinct community configuration rather than a simple return to the control state. Thus, D279 may reshape the microbial community under inflammatory conditions, rather than completely normalizing the microbiota to its pre-challenge configuration.

Changes in the selected bacterial taxonomic groups were also observed. However, broad taxonomic categories, such as Clostridia, encompass metabolically diverse organisms, and relative abundance alone cannot be directly equated with specific metabolic functions [33]. Therefore, the present microbiota data indicate an association between D279 supplementation and altered community structure. However, they are insufficient for the identification of individual bacterial taxa as the direct source of the observed SCFA changes.

Consistent with this interpretation, LPS was associated with a reduction in the contents of selected SCFAs, including n-valeric and n-butyric acids, whereas D279 partially attenuated these reductions. SCFAs contribute to intestinal homeostasis by affecting epithelial integrity, immune regulation, and mucosal metabolism [34,35]. Therefore, the preservation of selected SCFAs may represent an additional component of the intestinal environment associated with D279-mediated protection. Nevertheless, the present findings do not establish whether microbiota remodeling or SCFA preservation is causally involved in the protective effect of D279. Rather, the microbiota and SCFA results complement the epithelial, mucus, and immune findings. These findings suggest that D279 affects intestinal resilience through coordinated changes across multiple biological aspects [36].

### 4.6. Postbiotic Interpretation, Limitations, and Future Directions

The present findings identify heat-killed D279 as a post-biotic candidate capable of promoting intestinal resilience during systemic inflammatory stress through coordinated effects on (i) innate immune responsiveness, (ii) epithelial barrier properties, (iii) mucus-associated defenses, and (iv) intestinal microbial and metabolic environments. Heat-killed bacterial preparations exhibit biological activity even when bacteria are no longer viable. Their bioactive structural components interact with immune and epithelial receptors in the host [6].

D279 affected macrophage responses, supported epithelial barrier properties in vitro, attenuated colonic structural and mucus-associated injury in vivo, and was associated with changes in the microbiota and selected SCFA profiles. The distinct microbial configuration observed after D279 supplementation further suggests that protection was not necessarily linked to the complete restoration of an unchallenged intestinal state. Rather, it was likely linked to adaptive remodeling of the intestinal environment under inflammatory stress [37]. Figure 9 shows a schematic summary of the observed effects and proposed mechanisms underlying D279-mediated intestinal protection.

**Figure 9.**
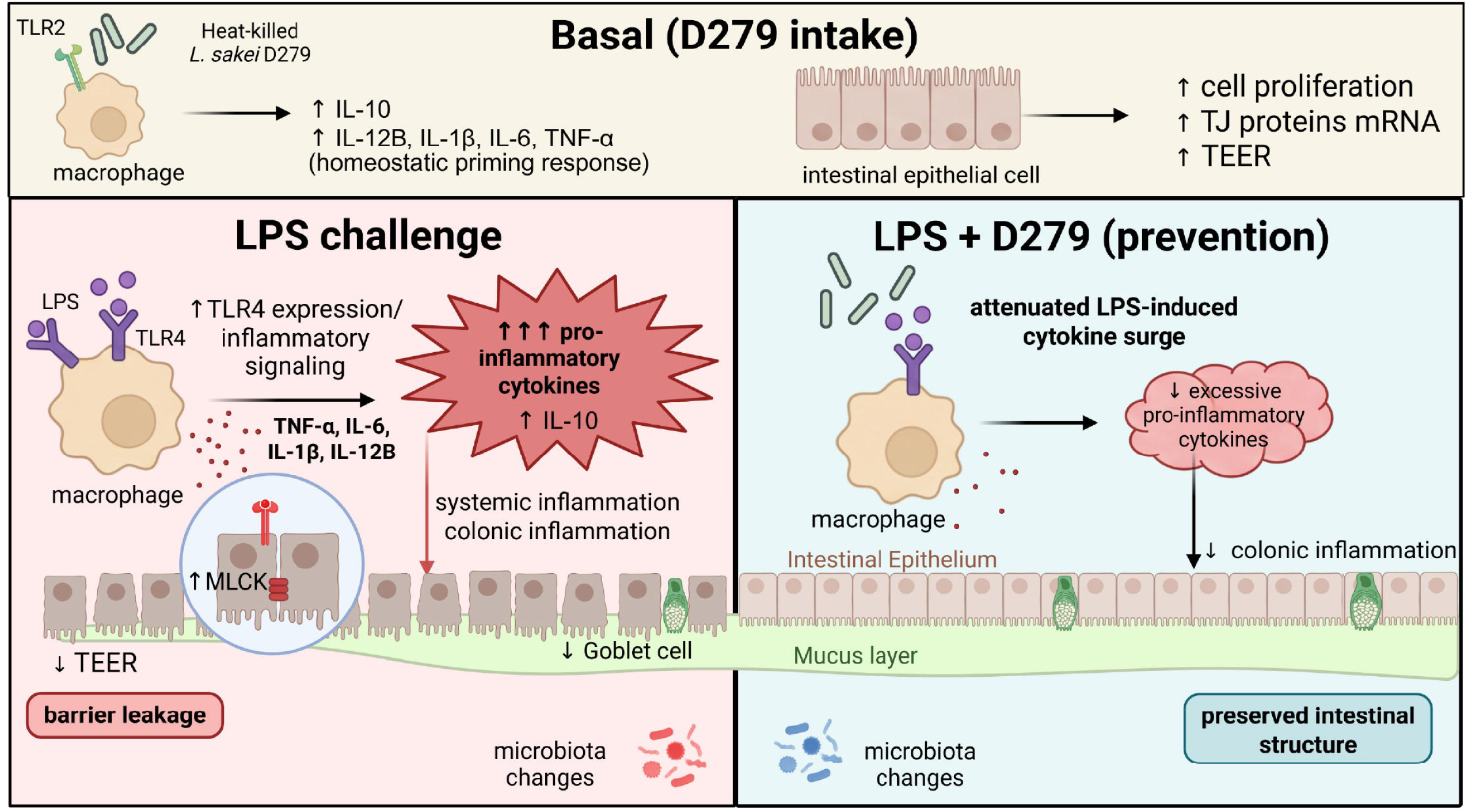
**Proposed mechanism underlying the protective effects of heat-killed *L. sakei* D279 against LPS-induced intestinal inflammation and barrier dysfunction.** The schematic summarizes the effects of D279 under basal conditions and during systemic LPS-induced inflammatory stress. Under basal conditions, the oral administration of heat-killed *L. sakei* D279 may promote intestinal homeostasis by stimulating macrophage-derived IL-10 and enhancing intestinal epithelial cell proliferation, expression of genes encoding TJ proteins, and TEER. In contrast, systemic LPS challenge activates TLR4-mediated inflammatory signaling, resulting in excessive pro-inflammatory cytokine production, macrophage activation, intestinal barrier disruption, goblet cell loss, and colonic inflammation. D279 supplementation attenuates LPS-induced inflammatory responses, preserves intestinal epithelial architecture and goblet cell abundance, and maintains the mucus layer, thereby alleviating intestinal barrier damage and colonic inflammation. These findings suggest that heat-killed D279 cells support intestinal homeostasis and protect against inflammation-associated intestinal injury. Illustrations were created with BioRender.com. LPS, lipopolysaccharide; TEER, transepithelial electrical resistance

The biological activity of D279 has been demonstrated in several contexts. In a previous study, oral administration of pasteurized D279 protected mice against influenza virus infection. This protective effect was characterized by enhanced natural killer cell activity and reduced viral load in the lungs. D279 also induced IL-12 production and preferentially stimulated IFN-λ expression in vitro, suggesting that its immunomodulatory activity extends beyond the intestinal inflammatory setting investigated in the present study [11]. Together with the present findings, these observations indicate that D279 exhibits biological activity in distinct experimental models involving innate immune regulation. However, whether common molecular mechanisms underlie these effects on influenza infection and intestinal inflammatory injury remains unclear.

This study had several limitations. First, the intraperitoneal LPS model represents acute systemic inflammation and does not fully reproduce chronic or multifactorial human intestinal injury. Second, although colonic inflammatory responses were evaluated, the causal relationships between specific inflammatory mediators and barrier protection were not experimentally isolated. Third, TJ regulation was largely assessed at the mRNA level; TJ protein localization and MLCK activity or phosphorylation were not directly quantified. Fourth, the present microbiota and SCFA findings only demonstrate associations. Therefore, functional experiments are required to determine whether these changes directly contribute to D279-mediated protection.

Future studies incorporating intestinal permeability assays, TJ protein localization, MLCK signaling analyses, and microbiota-function interventions (e.g., SCFA supplementation or microbiota depletion/manipulation approaches) are required to clarify the relative contributions of epithelial, immune, mucus, and microbial mechanisms to D279-mediated protection. In addition, a clinical assessment of heat-killed D279 is required to determine the translational relevance and verify whether these mechanistic signatures track clinically meaningful benefits. Specifically, future clinical investigations should use endpoints related to gastrointestinal symptoms, intestinal barrier function, fecal microbiota composition, SCFA profiles, and markers of intestinal inflammation.

## 5. Conclusions

Oral administration of heat-killed *L. sakei* D279 attenuated systemic LPS-induced colonic injury and inflammatory responses in mice. This strain preserved the colon length and mucosal architecture, mitigated goblet cell loss, and attenuated LPS-associated upregulation of *MLCK* expression, indicating its role in the maintenance of intestinal barrier homeostasis. Consistent with these in vivo findings, D279 enhanced the epithelial barrier function and reduced LPS-induced barrier impairment in IEC-6 cells. Ex vivo macrophage experiments further showed that D279 modulates innate immune responses and suppresses LPS-driven inflammatory responses, suggesting that immunomodulation contributes to its intestinal protective effects.

D279 supplementation was also associated with partial modification of LPS-induced changes in the intestinal microbiota and selected fecal SCFAs. Although causal relationships among immune regulation, epithelial and mucus barrier preservation, and microbiota/metabolite changes have not been established, these findings suggest that the protective activity of heat-killed D279 involves coordinated effects across multiple intestinal compartments.

Collectively, the present findings identify heat-killed *L. sakei* D279 as a promising postbiotic candidate for the promotion of intestinal resilience under systemic inflammatory stress. Further studies, including clinical investigations, are required to determine whether these protective effects can be translated into human intestinal health.

## Acknowledgements

The authors wish to thank the Tokushima Molecular Pathology Institute, Inc. for technical assistance with histological staining and slide preparation. We also acknowledge support from the TechnoSuruga Laboratory for SCFA analyses, gut microbiota sequencing/analysis, and microbial DNA extraction. Plasma biochemical analyses were conducted by Oriental Yeast Co., Ltd. We would also like to thank Editage (www.editage.jp) for English language editing. We further thank our colleagues at the Niigata Laboratory, MGC, for their helpful discussions and advice throughout the study.

## Ethical considerations

The animal experiments were approved by the Animal Care and Experimentation Facility Committee of the Mitsubishi Gas Chemical Company, Niigata Research Laboratory (Approval No. NA26-01).

The study was conducted in accordance with the committee’s rules and the ARRIVE guidelines.

## Declaration of generative AI and AI-assisted technologies in the manuscript preparation process

During the preparation of this work, the authors used Exabase in order to generate HTML code for a custom image analysis application used to quantify Alcian blue–positive area within regions of interest. After using this tool, the authors reviewed and edited the content as needed and take full responsibility for the content of the published article.

