## Supplementary material for "Heat-killed Latilactobacillus sakei D279 Attenuates Lipopolysaccharide-Induced Intestinal Inflammation and Barrier Disruption": Suplementary data

**Supplementary Materials**

To ensure reproducibility, all assay kits and antibodies used in this study are listed in Supplementary Table S1, including supplier information and catalog numbers.

**Supplementary Table S1. Assay kits, reagents, and antibodies used in this study**

| **Category** | **Item** | **Supplier** | **Catalog No.** | **Application** |
| --- | --- | --- | --- | --- |
| Assay kit | Cell Counting Kit-8 (CCK-8) | Dojindo | CK04 | Cell viability |
| Assay reagent | CellROX Green | Thermo Fisher Scientific | C10444 | ROS detection |
| ELISA kit | Mouse IL-12 ELISA kit | Proteintec | KE10014 | Cytokine quantification |
| ELISA kit | Lipoteichoic acid (LTA) ELISA kit | Cell Biolabs, Inc. | AKR-5153 | LTA quantification |
| Antibody | Anti-NF-κB p65 | Proteintech | 10745-1-AP | Immunofluorescence (IF) |
| Antibody | CoraLite488 Donkey Anti-Rabbit IgG | Proteintech | SA00013-6 | Immunofluorescence (IF) |
| Antibody | CoraLite®594-conjugated ZO-1 Polyclonal antibody | Proteintech | CL594-21773 | Immunofluorescence (IF) |
| Antibody | Claudin 1 Polyclonal antibody unconjugated | Proteintech | 13050-1-AP | Immunofluorescence (IF) |
| Antibody | Anti-TLR2 | Abeomics | 10-3026-25 | Blocking assay |
| Antibody | Anti-TLR4 | Proteintech | 66350-1-1G | Blocking assay |
| Antibody | Anti-TLR9 | Abeomics | 10-3003-25 | Blocking assay |
| Antibody | Anti-Dectin-1 | Abcam | Ab37154 | Blocking assay |
| RNA extraction kit | [Kit name] | Quick-RNA Miniprep Plus Kit | R1057 | RNA extraction |

**Supplementary Results**

**Cellular Association of FITC-Labeled D279 with Macrophages**

To determine whether D279 physically interacts with macrophages ex vivo, FITC-labeled D279 was incubated with macrophages and examined by fluorescence microscopy. This experiment was performed as a preliminary assessment of whether labeled D279 could be detected in association with cells under the experimental conditions.

Macrophage-like cells were seeded in 96-well plates at 1.0 × 10⁵ cells/mL and cultured for 24 h. FITC-labeled D279 was then added at a final concentration of 10 μg/mL, followed by an additional 24-h incubation. Bright-field and fluorescence images were obtained after incubation.

In the absence of D279, macrophage-like cells were observed in bright-field images, with no apparent D279-derived fluorescence. In contrast, cells incubated with FITC-D279 showed distinct fluorescence signals associated with the cells. The enlarged images further demonstrated cell-associated fluorescent signals.

These observations confirmed that FITC-labeled D279 could be detected in association with macrophage-like cells using fluorescence microscopy. However, because conventional fluorescence microscopy does not distinguish bacterial attachment to the cell surface from intracellular localization, these data demonstrate cellular associations rather than definitive intracellular uptake.

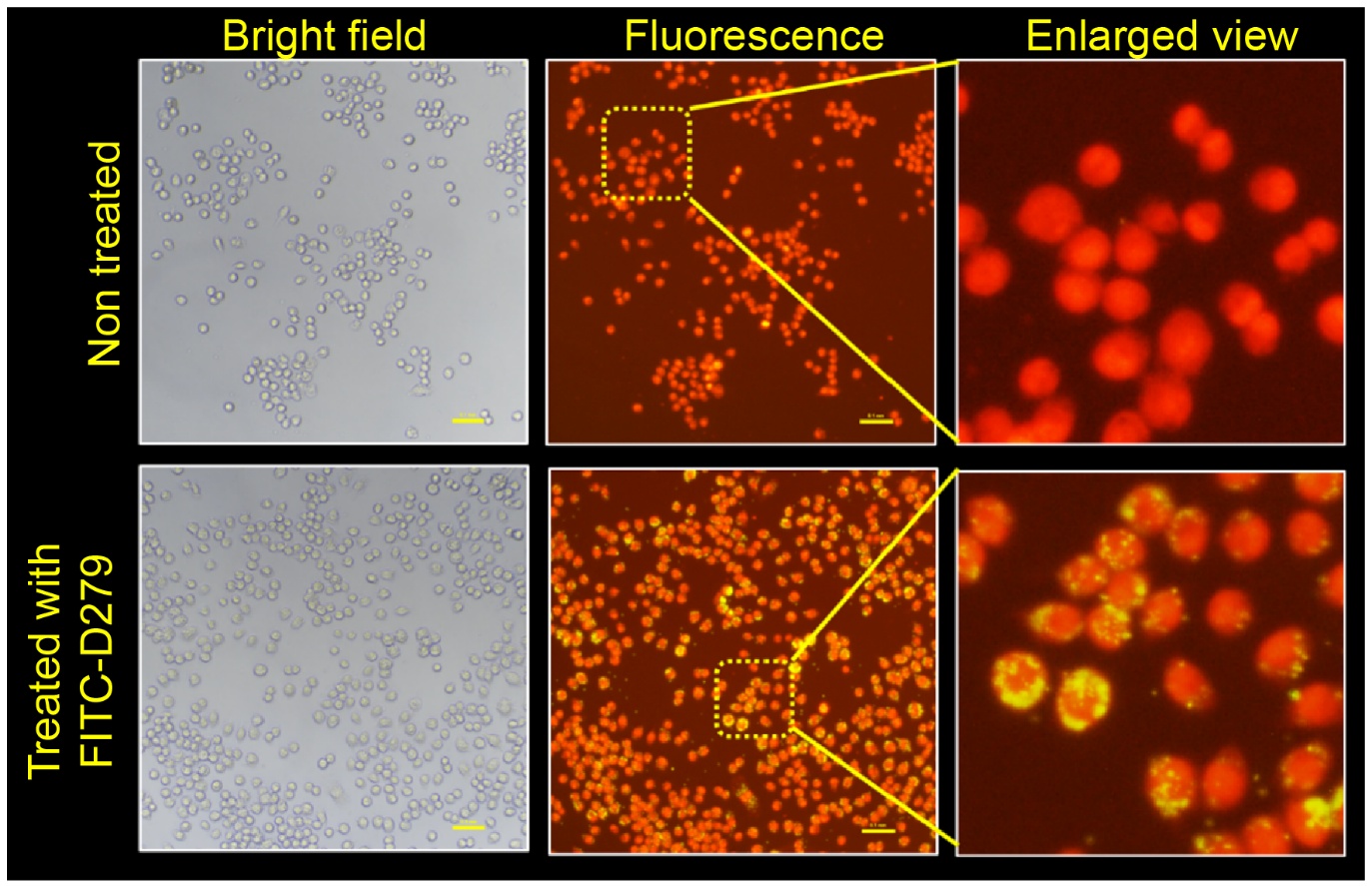

**Supplementary Figure S1. Cellular association of FITC-labeled D279 with macrophage-like cells.**
Macrophage-like cells were incubated with FITC-labeled D279 (10 μg/mL) for 24 h. Representative bright-field and fluorescence images are shown for cells without D279 and those treated with FITC-D279. Enlarged views of the indicated regions are shown on the right. Scale bar; 100 μm.

**Contribution of pattern-recognition receptors to D279-induced IL-12p70 production**

To investigate the receptor pathways involved in the macrophage response to D279, the contributions of several pattern-recognition receptors were examined using receptor-blocking antibodies in an ex vivo macrophage assay. Macrophages were treated with antibodies against TLR2, TLR4, TLR9, or Dectin-1 before stimulation with D279. IL-12p70 production was subsequently measured.

Inhibition of TLR2 or Dectin-1 markedly reduced IL-12p70 production compared to that in the mock-treated conditions. In contrast, the inhibition of TLR4 or TLR9 resulted in a comparatively minimal reduction in IL-12p70 production. These findings indicate that TLR2- and Dectin-1-dependent pathways contribute to the macrophage response to D279, whereas TLR4 and TLR9 may play less prominent roles under these experimental conditions. Because TLR2 and Dectin-1 recognize the structural characteristics of microbial cell surfaces, these findings further demonstrate the involvement of cell wall-associated components in D279-mediated macrophage activation. However, receptor blockade does not identify specific bacterial ligands responsible for receptor activation. Thus, these findings confirm the involvement of TLR2/Dectin-1-associated recognition pathways but do not establish a particular cell wall component as an active immunostimulatory factor.

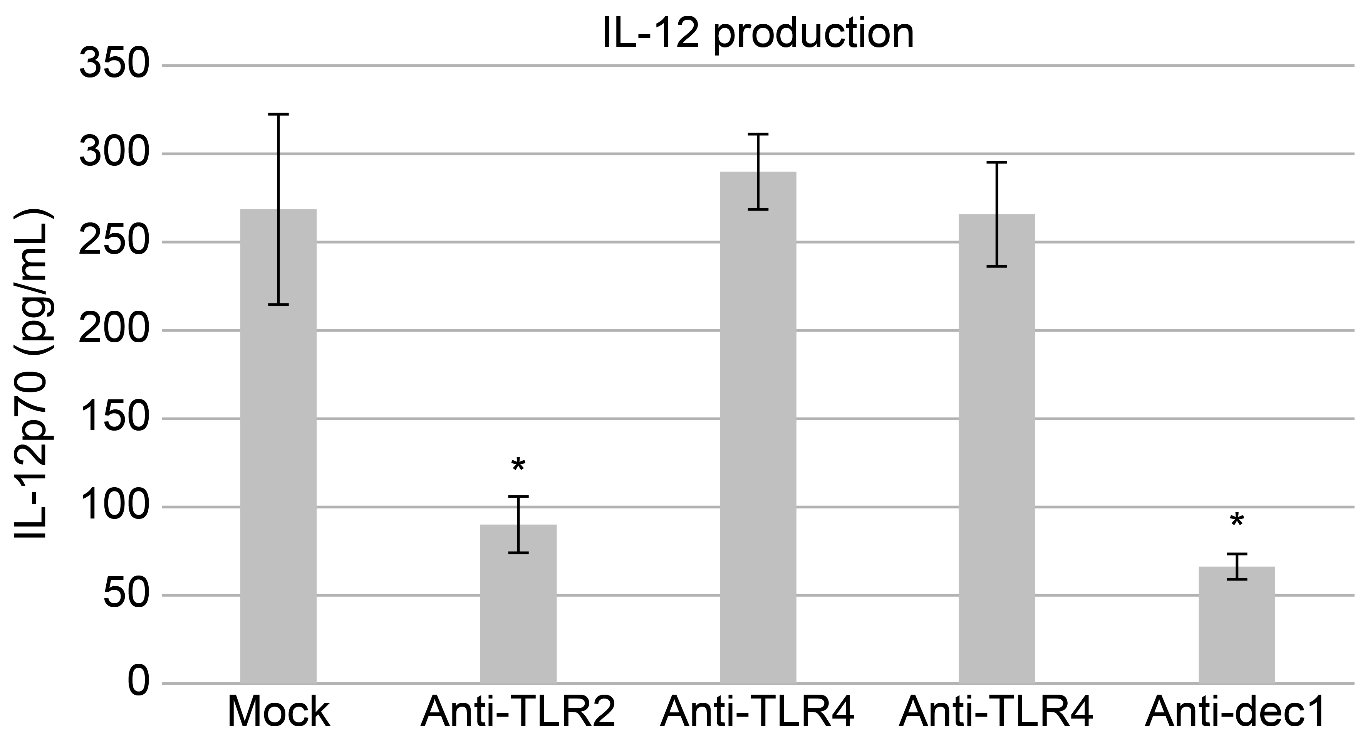

**Supplementary Figure S2. Contribution of pattern-recognition receptors to D279-induced IL-12p70 production.**
IL-12p70 production, as measured in ex vivo macrophages treated with antibodies against TLR2, TLR4, TLR9, or Dectin-1 before stimulation with D279.

**Lipoteichoic acid content in D279 and comparator lactic acid bacterial strains**

As receptor-blocking experiments suggested the involvement of cell wall-associated recognition pathways, we examined the abundance of lipoteichoic acid (LTA), a major cell envelope component of gram-positive bacteria and a potential contributor to host immune recognition.

LTA in whole bacterial preparations was quantified using a commercial LTA ELISA kit. Briefly, 100 mg of bacterial powder was suspended in 1 mL of PBS, sonicated, incubated at 4 °C, and centrifuged 10,000 × *g*. The resulting supernatant was used for ELISA. LTA levels were compared between D279, the *Lactobacillus sakei* type strain, and other lactic acid bacterial strains.

D279 showed an LTA level of approximately 6,600 ng/mL—comparable to that of the *L. sakei* type strain (approximately 6,800 ng/mL). In contrast, LAB A, B, C, and D showed lower LTA levels, ranging from approximately 3,500 to 4,800 ng/mL. Thus, among the strains examined, D279 contained a relatively high level of LTA, comparable to that of the *L. sakei* type strain. These findings suggest that LTA contributes to the immunostimulatory properties of D279. However, the present data demonstrate an association between LTA abundance and bacterial strain phenotype; they do not establish LTA as the definitive active component. Therefore, further experiments using preparations with depleted or purified LTA preparations are required to determine its direct contributions to D279-mediated immune activation.

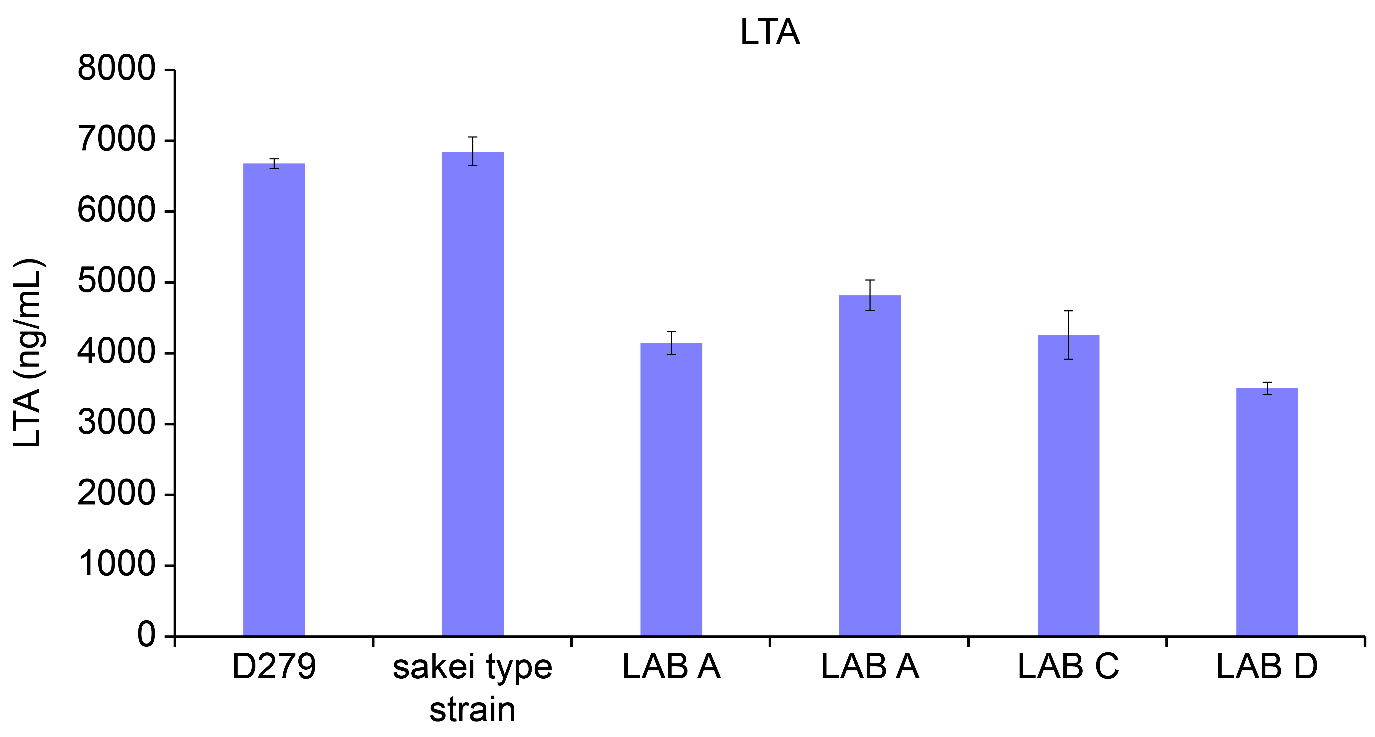

**Supplementary Figure S3. LTA content of D279 and comparator lactic acid bacterial strains.**
LTA levels, as quantified from whole bacterial preparations of D279, the *L. sakei* type strain, and LAB A–D. Data are presented as the mean ± SD.

**Effects of long-term D279 supplementation on the expression of genes related to the intestinal barrier in healthy mice**

D279 was selected based on its strong IL-12-inducing activity in an ex vivo mouse splenocyte assay. However, its effects under physiological conditions have not been extensively investigated. RNA samples obtained from a previous feeding study were analyzed to determine whether long-term D279 supplementation affects the expression of intestinal barrier-associated genes in healthy animals.

The mice were maintained under non-stress conditions and received either a standard diet or the same diet supplemented with 0.1% (w/w) D279 for 8 weeks. Total RNA was isolated from the small intestine, and the mRNA levels of the TJ–associated genes *occludin, ZO-1,* and *claudin-1* were quantified. Dietary D279 significantly upregulated the expression of these transcripts by approximately 1.3- to 1.8-fold compared to that in the control group. These findings suggest that long-term D279 intake promotes the expression of genes involved in intestinal epithelial barrier maintenance, even in the absence of overt inflammatory stimulation.

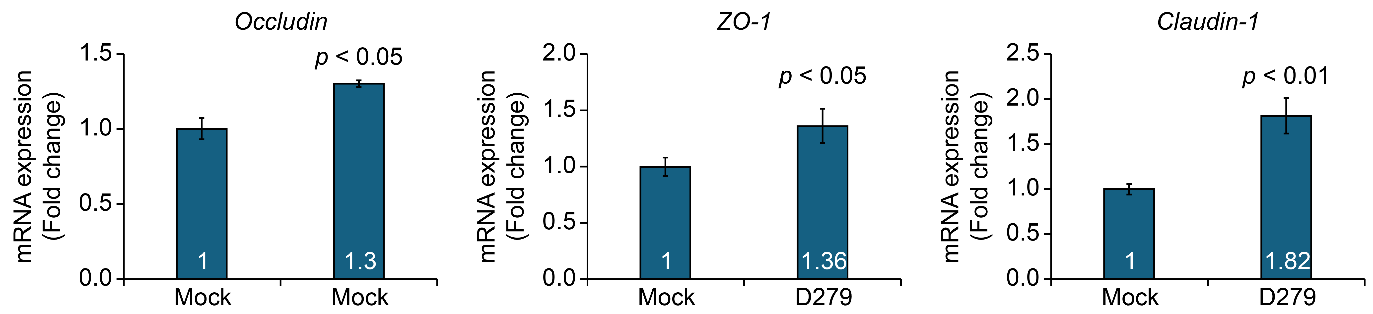

**Supplementary Figure S4. Effects of long-term D279 supplementation on the expression of intestinal barrier-related genes in healthy mice.**
After the feeding period, small-intestinal mRNA levels of *Ocln*, *Zo1*, and *Cldn1*, as quantified by RT–qPCR in mice fed either a standard control diet or the same diet supplemented with 0.1% (w/w) D279 for 8 weeks. Data are expressed as the mean ± SE.
